# Signatures of recent positive selection in enhancers across 41 human tissues

**DOI:** 10.1101/534461

**Authors:** Jiyun M. Moon, John A. Capra, Patrick Abbot, Antonis Rokas

## Abstract

Evolutionary changes in enhancers are widely associated with variation in human traits and diseases. However, studies comprehensively quantifying levels of selection on enhancers at multiple evolutionary time points during recent human evolution and how enhancer evolution varies across human tissues are lacking. To address these questions, we integrated a dataset of 41,561 transcribed enhancers active in 41 different human tissues (FANTOM Consortium) with whole genome sequences of 1,668 individuals from the African, Asian, and European populations (1000 Genomes Project). Our analyses based on four different metrics (Tajima’s *D*, *F*_ST_, H12, *nS*_L_) showed that ~5.90% of enhancers considered showed evidence of recent positive selection and that genes associated with enhancers under positive selection are enriched for diverse immune-related functions. The distributions of these metrics for brain and testis enhancers were often statistically significantly different compared to those of other tissues; the same was true for brain and testis enhancers that are tissue-specific compared to those that are tissue-broad and for testis enhancers associated with tissue-enriched and non-tissue-enriched genes. These differences varied considerably across metrics and tissues and were generally due to changes in distributions’ shapes rather than shifts in their values. These results suggest that many human enhancers experienced recent positive selection throughout multiple time periods in human evolutionary history, that this selection occurred in a tissue-dependent and immune-related functional context, and that much like the evolution of their coding counterparts, the evolution of brain and testis enhancers has been markedly different from that of enhancers in other tissues.

## Introduction

Enhancers are *cis*-acting DNA segments that, either independently of or in concert with other regulatory elements, control spatial, temporal and quantitative aspects of gene expression (Ong and Corces 2011; Rubinstein and de Souza 2013; Long et al. 2016). While the precise architecture of enhancers is still debated (Long et al. 2016), a typical enhancer contains multiple transcription factor binding sites (TFBS) arranged in specific order and distance from one another. Enhancers facilitate initiation of gene transcription by helping to recruit RNA Polymerase II, general transcription factors, and additional components of the transcriptional machinery to the gene’s promoter (Pennacchio et al. 2013; Rubinstein and de Souza 2013). Multiple enhancers, each with its own repertoire of TFBS, can regulate the activities of a gene in a tissue-specific manner or across distinct developmental stages, enabling enhancers to alter its expression patterns in a particular context without affecting expression of other genes (Wray 2007; Sholtis and Noonan 2010). Enhancers in the human genome are more numerous than protein-coding genes (Pennacchio et al. 2013), facilitating the induction of diverse gene expression programs in different spatial and temporal contexts (Long et al. 2016).

Changes in gene regulation have long been thought to play a major role in the adaptive evolution of human traits (King and Wilson 1975; Carroll 2005). One reason for regulatory regions in general, and enhancers in particular, as preferential targets of selection is that, compared to protein-coding regions, they tend to be modularly organized (Sholtis and Noonan 2010). This modular organization means that mutations in enhancers are less likely to have pleiotropic effects and more likely to contribute to phenotypic evolution (Carroll 2005; Wray 2007; Rebeiz and Tsiantis 2017). In the context of human evolution, several studies suggest that evolutionary changes in enhancers might have played a major role in the acquisition of human-specific traits. For example, a considerable proportion of regions that has experienced accelerated evolution in the human lineage are developmental enhancers active in the brain (Capra et al. 2013), and *cis*-regulatory regions of genes with neurological and nutritional roles have been shown to exhibit evidence of accelerated evolution in the human lineage (Rockman et al. 2005; Haygood et al. 2007). Accelerated evolution is one signature of positive selection, and several studies suggest that recent selection might also have preferentially acted on human *cis*-regulatory regions (Wray 2007). For instance, Rockman et al. found that the promoter region of a gene that encodes the precursor of an opioid neuropeptide (*PDYN*) exhibits a significant degree of population differentiation between human populations, especially between Europeans and East Asians, which is suggestive of local adaptation (Rockman et al. 2005). Similarly, patterns of variation in the promoter region of the *LCT* gene that confers lactase persistence in Africans are consistent with the action of selective sweeps (Tishkoff et al. 2006). Finally, SNPs with evidence of recent positive selection are more likely to be associated with expression of nearby genes than random SNPs, and this enrichment is strongest for Yorubans (Kudaravalli et al. 2008).

More broadly, genome-wide studies have found that, compared to protein-coding regions, enhancers are enriched for variants that are statistically associated with various human diseases (e.g., inflammatory diseases, metabolic diseases) (Andersson et al. 2014; Karnuta and Scacheri 2018). Moreover, recent studies have argued that human population-level differences in transcriptional responses to infection likely resulted from local adaptation, further supporting the idea that selection on enhancers has contributed to recent human evolution (Nédélec et al. 2016). In addition, multiple events, such as migrations and shifts in cultural practices, have occurred in different time periods during recent human evolution (Karlsson et al. 2014), likely introducing novel selective agents. Given the role of enhancers in contributing to phenotypic differences, it is likely that enhancers experienced selection events in response to such selective pressures.

Each human tissue serves a particular physiological function so it is reasonable to hypothesize that the genetic elements active in each tissue are influenced by distinct selective pressures that shape their evolutionary rates (Wray 2007; Gu and Su 2007). For example, both gene expression and protein-coding sequence divergence patterns might be expected to be more conserved in developmentally constrained tissues (e.g., nervous tissues) than in developmentally relaxed tissues (e.g., testis or endocrine tissues, such as pancreas). Alternatively, certain functions of tissues (e.g., reproductive processes for testis) might have experienced increased levels of positive selection (Khaitovich 2005). Early studies examining patterns of divergence in gene expression and protein-coding sequence evolution among mammals found that both were lowest for nervous tissues (e.g., brain, cerebellum) and greatest for testis (Wray 2007; Khaitovich 2005; Brawand et al. 2011). More recent examination of a larger number of human tissues has provided further support for this hypothesis. For example, the correlation between expression levels of genes and the strength of purifying selection (as assessed by *d*N/*d*S) is strongest for the brain, while this correlation is lowest for liver, placenta and testis (Kryuchkova-Mostacci and Robinson-Rechavi 2015).

Hitherto, studies investigating the variation of evolutionary patterns among tissues have mainly focused on inter-species divergence of gene expression and of protein-coding sequences. However, studies comprehensively quantifying levels of selection on enhancers at multiple evolutionary time periods during recent human evolution and how enhancer evolution varies across human tissues are lacking. Furthermore, it is plausible, if not likely, that distinct selective pressures are also acting on regulatory elements in different tissues, further contributing to the global differences in gene expression patterns among the tissues described above (Ong and Corces 2011; Rubinstein and de Souza 2013).

To examine the influence of selection on enhancers at multiple evolutionary time periods during recent human evolution and how enhancer evolution varies across human tissues, we used a dataset of 41,561 enhancers active in 41 human tissues and genotype data of 1,668 individuals from three different super-populations from the 1000 Genomes Project to calculate signatures of recent selection that happened at different time points and via different modes (i.e., hard vs soft selective sweeps). We found that on average, 5.90% of enhancers exhibit evidence of recent positive selection and that their putative target genes are enriched for immunity-related functions. Furthermore, we found that enhancers expressed in the testis and brain exhibit statistically different patterns of recent evolution compared to numerous other tissues. Examination of patterns of recent evolution between enhancers that are active in only one tissue and those active in two or more tissues revealed statistically significant differences for several tissues, including brain and testis; we found similar results when we studied enhancers associated with tissue-enriched and non-tissue-enriched genes in these two tissues. Our results suggest that human enhancers, in particular ones associated with immune-related genes, have experienced different modes of recent positive selection at different periods of recent human evolution. Furthermore, patterns of selection on human enhancers differ between tissues, including for brain and testis, a pattern reminiscent of their protein-coding counterparts.

## Methods

### Genetic Variation Data

To examine signatures of natural selection on human enhancers, we used the whole genome sequence data for 1,668 individuals from Phase 3 of the 1000 Genomes Project (The 1000 Genomes Project Consortium et al. 2015) (for information on data sources, see File S1). The 1,668 individuals represent three major human populations (661 Africans, 503 Europeans, and 504 East Asians). We used these three populations because their demographic histories have been most confidently estimated and excluded the other two (Admixed Americans and South Asians) because their demographic histories are more complex.

### FANTOM Enhancer Data

To study patterns of recent evolution on human enhancers active in different tissues, we used the enhancers detected in 41 human tissues compiled by the Functional Annotation of the Mammalian Genomes (FANTOM) project (Figure 1 & File S1) (Andersson et al. 2014) based bidirectional transcription identified via cap analysis of gene expression (CAGE) in diverse human cell and tissue types (The FANTOM Consortium et al. 2014). We further classified the enhancers into ‘tissue-specific’ enhancers that are detected in only a single tissue, and ‘tissue-broad’ enhancers that are detected in two or more tissues. As genomic regions located on sex chromosomes exhibit patterns of genetic variation that differ from those observed in autosomal genomic regions (Schaffner 2004), we restricted our analyses to only autosomal enhancers.

**Figure 1.**
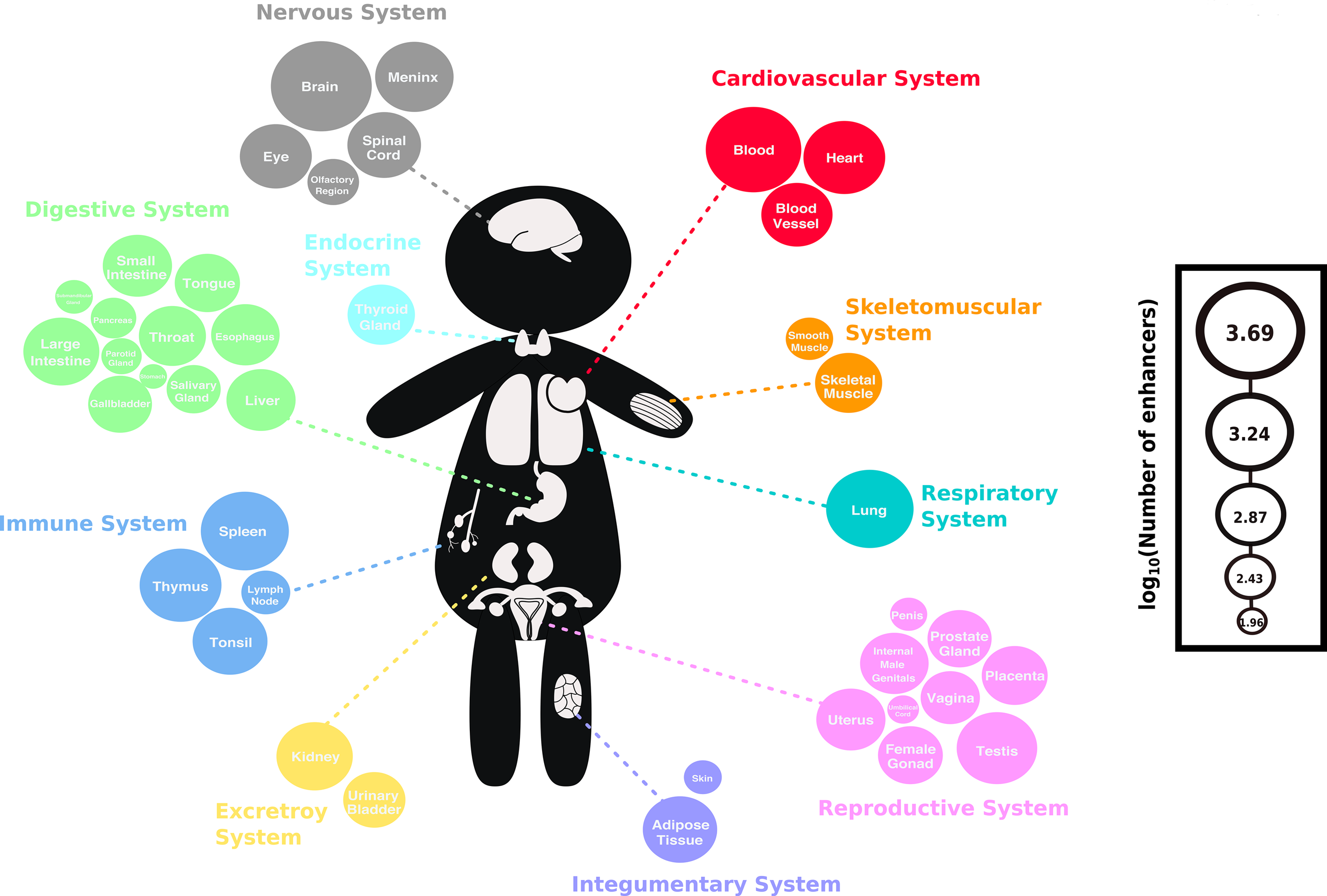
Visual summary of the FANTOM5 enhancer data set used in this study. Each circle represents a tissue and all 41 tissues are grouped into non-overlapping organ systems and are color-coded accordingly. The size of the circle is proportional to the log_10_ of the number of enhancers active within each tissue (Table S1: from highest to lowest: brain: 4,883, blood: 3,657, spleen: 2,429, lung: 2,366, thymus: 1,741, heart: 1,737, testis: 1,621, meninx: 1,447, kidney: 1,294, large-intestine: 1,266, tonsil: 1,193, adipose-tissue: 1,161, spinal-cord: 1,115, eye: 1,019, blood-vessel: 982, small-intestine: 880, uterus: 877, liver: 875, internal-male-genitalia: 846, esophagus: 845, thyroid-gland: 795, skeletal-muscle: 784, throat: 782, placenta: 735, tongue: 708, female-gonad: 693, prostate-gland: 667, gallbladder: 654, urinary-bladder: 602, vagina: 512, salivary-gland: 387, olfactory-region: 338, lymph-node: 292, smooth-muscle: 270, pancreas: 243, parotid-gland: 184, skin: 161, submandibular-gland: 159, penis: 154, umbilical-cord: 115, stomach: 92).

### Human Protein Atlas Data

To compare the patterns of recent evolution between enhancers associated with genes that exhibit higher expression levels in one tissue compared to others and those that are not in our tissues of interest, the brain and testis, we downloaded the list of tissue-enriched genes for the brain and testis tissues from the Human Protein Atlas database (Uhlen et al. 2015) (version 88; download date: January 17^th^, 2018): the Human Protein Atlas Database defines a gene as tissue-enriched if its mRNA levels in a given tissue are at least five-fold higher compared to its levels of expression in all other tissues. We defined ‘non-tissue-enriched’ genes as those genes that are not labeled as ‘tissue-enriched’ according the to the Protein Atlas database.

### Identifying Signatures of Recent Positive Selection

To identify signatures of recent positive selection on human enhancers, we calculated four metrics aimed to capture genetic signatures left from the action of selection at different time ranges during recent human evolution (Sabeti et al. 2006; Vitti et al. 2013; Moon et al. 2018) using previously described methodology (Moon et al. 2018). Briefly, the four metrics that we used can be divided into two categories: those that are designed to detect hard selective sweeps (Tajima’s *D*, Weir & Cockerham’s *F*_ST_ and *nS*_L_) and those designed to detect soft selective sweeps (H12). Furthermore, the three tests for hard selective sweeps are most sensitive to selection events that occurred at different time points in human history: Tajima’s *D* can detect selection that happened approximately 250,000 - 200,000 years ago; *F*_ST_ can identify signatures of local selection that occurred following the out-of-African migration approximately 75,000 - 50,000 years ago; finally, *nS*_L_ has power to detect signatures left by selection events that occurred approximately 20,000 - 10,000 years ago (Sabeti et al. 2006; Moon et al. 2018). Following the methods of Moon et al. (Moon et al. 2018) for each enhancer region in every tissue we calculated Tajima’s *D* using the R package *PopGenome*, version 2.1.6 (Pfeifer et al. 2014) and the weighted Weir & Cockerham’s fixation index (*F*_ST_) (Weir & Cockerham 1984) among all three populations using *VCFtools*, version 0.1.13 (Danecek et al. 2011). In addition, we calculated the *nS*_L_ statistic (number of segregating sites by length; (Ferrer-Admetlla et al. 2014)) using *Selscan*, version 1.2.0 (Szpiech and Hernandez 2014). For each enhancer, we calculated *nS*_L_ for the region extending 50 kilobases (kb) upstream and downstream using the default settings of *Selscan*, except for the minor allele frequency (MAF) cut-off value, for which we used 0.01. We calculated this metric for each of the three populations, as haplotype-based methods are known to detect selection events that occurred after the out-of-African migration (Voight et al. 2006; Sabeti et al. 2006). We used the maximum absolute un-standardized *nS*_L_ values calculated over the entire window to represent each enhancer region. Finally, for each enhancer region we also calculated the H12 index using a custom script, based on Garud *et al.*’s original script (Garud et al. 2015). The values of all enhancers in the 41 human tissues for all metrics can be found in the Figshare repository.

To test for statistical differences in the distributions of each of the four metrics between any two tissues, we also carried out pairwise Kolmogorov-Smirnov tests (two-sided). Visualization of the data (Figure S1) revealed that assumptions of normality and equal variance could not be upheld, and therefore, we chose the non-parametric Kolmogorov-Smirnov test which makes no assumptions regarding the shape of the distributions and carried out this test in the R programming environment. We also conducted this test to examine whether there are any significant differences between the distributions of patterns of recent evolution between tissue-specific and tissue-broad enhancers, as well as between enhancers that are associated with tissue-enriched and non-tissue-enriched genes. All statistical tests were followed by *post hoc* Bonferroni corrections in R. The results of the statistical tests for all metrics can be found in the Figshare repository.

### Simulations of Neutral Evolution

The action of non-adaptive processes and demographic changes, such as genetic drift and population expansion, can produce patterns of variation that these tests may mistake for positive selection (Sabeti et al. 2006). To determine the likelihood of the empirical values being generated by the action of selection, we carried out simulations of neutral evolution based on a model of recent human demographic history and compared the observed values to expected values under neutrality. Simulations of neutrality were conducted using *SLiM*, version 2.4.1 (Messer 2013; Haller and Messer 2017). Following Gravel et al., we used previously calculated demographic parameters for the three populations included in our study (Gravel et al. 2011; Messer 2013). Detailed information on the parameters used for the simulations can be found in the Supplementary File S1.

We simulated the genotypes of 661, 503, and 504 individuals corresponding to the numbers of individuals from each of the three populations analyzed. Since there are 41,561 autosomal enhancers in the FANTOM dataset and anywhere from 92 (stomach) to 4,883 (brain) enhancers in any given tissue (Table S1), carrying out individual simulations for each enhancer was computationally prohibitive. To reduce the computational load of our simulations, we focused on simulating the mutational profile of an average enhancer for a given tissue under a model of neutral evolution. Specifically, to carry out the neutral simulations for enhancers in a given tissue, we used the average length of all the autosomal enhancers found in that tissue (these values can be found in the Figshare repository) and a fixed recombination rate of 1×10^−8^. For each tissue, we carried out 10,000 simulations. We next used these simulated sequences to calculate Tajima’s *D* values, Weir & Cockerham’s *F*_ST_, and H12 as described above. We calculated an empirical *p*-value for a given enhancer for any metric (including for *nS*_L_, the simulation process of which is described below) as the proportion of simulated sequences that obtained scores equal to or more extreme than the observed value. We used a *p*-value of 0.05 as cutoff for significance; enhancers with *p*-values lower than 0.05 were considered to significantly deviate from neutral evolution and to have experienced selection.

### Recombination Rate Interpolation and Simulation of nS_L_ Values

As variation in recombination rates can affect the values of *nS*_L_ (Ferrer-Admetlla et al. 2014), we carried out a separate set of neutral simulations for the calculation of *nS*_L_ values by incorporating average recombination rates for each tissue. To do this, we used the genome-wide genetic map curated by the HapMap II consortium (Frazer et al. 2007), which provides pre-computed recombination rates (cM/Mb) for the variants included in the HapMap II project. More specifically, we carried out linear interpolation to infer the recombination rates of the variants in the 1000 Genomes Project dataset that are not included in the HapMap Phase II dataset. Next, we calculated the average recombination rate (cM/Mb) for the region spanning 50 kb up and downstream of each enhancer in a given tissue, and then used the average of all recombination rates of the enhancer regions in a given tissue as the recombination rate parameter (converted to probability of crossovers per bp) for the neutral simulations. The average recombination rates, as used in our *SLiM* simulations, for each tissue can be found in the Figshare repository. Using these calculated recombination rates, we then created 2,500 enhancers that span 50 kb upstream and downstream of the average length of all autosomal enhancers found in that given tissue and calculated *nS*_L_ values on these simulated enhancers using *Selscan* as described above. As with the actual enhancers, we only included variants with MAF greater than 0.01 in the calculations and used the maximum absolute un-standardized values to represent a given simulated window.

### Functional Enrichment Analyses and Semantic Similarity Calculations

To gain insight into the functions of genes associated with enhancers that show evidence of selection, we next carried out functional enrichment analyses. To associate enhancers with putative target genes, we used the transcription start site (TSS)-enhancer mapping file generated by Andersson et al. (Andersson et al. 2014) (file location information can be found in File S1), which was compiled by measuring the pairwise correlation between enhancer activity and transcription level of putative target genes. We carried out functional enrichment analyses on the list of these putative target genes using the R package *TopGO*, version 2.32.0 (Alexa and Rahnenfuhrer 2016). Detailed description of the files and commands used can be found in File S1. In short, we used the ‘weight’ algorithm that compares the significance scores of the connected nodes to explicitly account for the hierarchy of the gene ontology tree and carried out analyses for the three general ontologies: Biological Process (BP), Molecular Function (MF), and Cellular Component (CC). Subsequent corrections for multiple comparisons were carried out by calculating the FDR-adjusted *p*-values in the R statistical environment; only those GO terms with an FDR-adjusted *p*-value less than 0.05 were retained for further analyses. All *TopGO* enrichment analysis results for all the metrics can be found in the Figshare repository.

To quantify the overall similarity of the patterns of gene functional enrichment among different tissues, we calculated pairwise semantic similarity of the GO terms between any two tissues using a graph-based method (Wang et al. 2007), which takes into consideration the topology of the GO graph structure (i.e., the location of the GO terms in the graph and the relationship of the terms to their ancestor terms), as implemented in the R package *GOSemSim*, version 2.4.1 (Yu et al. 2010): this analysis was carried out for only those tissues that have 5 or more significant GO terms. To determine if the semantic similarity scores thus calculated significantly deviated from random expectations, we randomly sampled the same number of GO terms for each tissue from the pool of all GO terms associated with a given tissue 1,000 times and calculated the pairwise semantic similarity score; scores equal or greater than the 95^th^ percentile value of the distribution of semantic similarity scores were considered significant. The calculated semantic similarity values, along with the 95^th^ percentile values obtained from the 1,000 randomly sampled GO terms for each tissue pair, can be found in the Figshare repository.

### Transposable Element (TE) Data and Analyses

TE-derived sequences often contribute to the origin of new enhancers and the modification of regulatory networks (Lynch et al. 2015; Simonti et al. 2017). To determine if the enhancers with evidence of recent positive selection harbor higher proportions of TEs than those with nonsignificant deviations from neutral expectations, we used BEDOPS, version 2.4.35 (Neph et al. 2012) to overlap the enhancer regions with the RepeatMasker-annotated regions. We downloaded the RepeatMasker annotation track for the hg19 assembly from the UCSC Genome Browser (Kent et al. 2002; Casper et al. 2018; Raney et al. 2014). The location of the RepeatMasker annotation file and the specific commands used for BEDOPS can be found in Supplementary File S1. To test for statistical differences in the proportions of TEs between enhancers with evidence of recent selection and those that do not exhibit such significant deviation from neutral expectations, we carried out a 2×2 chi-squared test with Yate’s continuity correction using R. The results of the chi-squared tests can be found in the Figshare repository.

### GWAS Catalog Data Annotation

Previous studies have shown that many variants in human *cis*-regulatory regions are significantly associated with a broad range of complex traits and diseases (Lee and Young 2013; Andersson et al. 2014; Karnuta and Scacheri 2018). To investigate whether brain and testis enhancers that exhibit evidence of recent positive selection show enrichment of variants in the Genome-Wide Association Study (GWAS) catalog, we downloaded the NHGRI-EBI GWAS catalog, version1.0.2 (MacArthur et al. 2017) (download date: June 2^nd^, 2018) and first compared the SNPs that reside in the enhancer regions with GWAS catalog SNPs. We also included SNPs that are outside the enhancer regions but in complete linkage disequilibrium (LD) with the SNPs within the enhancer regions. We used Plink, version 1.0.9 (Chang et al. 2015) to obtain a list of SNPs that are in complete LD (r^2^ = 1.0) with those that lie within the enhancers of interest (‘complete LD SNPs’) and carried out the same analyses as described above. Detailed description of the Plink analyses can be found in Supplementary File S1. For subsequent analyses, we combined the GWAS hits of the SNPs that reside within the enhancers and those that are in complete LD with SNPs inside the enhancers and considered them together. We also carried out the same analyses for brain and testis enhancers that exhibit nonsignificant deviations from neutral expectations. To determine if there is a statistical difference in the enrichment of variants in the GWAS hits between enhancers that exhibit evidence of recent positive selection and those that do not, we carried out 2×2 chi-squared tests with Yate’s continuity correction using R. The full list of all the GWAS hits that overlap with all the variants considered, as well as the chi-squared test results, can be found in the Figshare repository.

## Results

### Enhancers experienced positive selection at different time ranges during recent human history

Calculation of four metrics of selection (Weir & Cockerham’s *F*_ST_, Tajima’s *D*, H12 and *nS*L) for all 41,561 enhancers in 41 tissues (these data can be found in the Figshare repository) revealed that an average 5.90% of enhancers have experienced recent positive selection considering all metrics and tissues (Figure 2). The proportions of enhancers that exhibit significant deviations from neutral expectations for each metric varied (Figure 2; Table S2). Specifically, greater fractions of enhancers show evidence of recent positive selection according to H12 and Tajima’s *D* metrics (H12: from 6.57% to 16.67%; Tajima’s *D*: 6.75% to 13.92%) than other metrics (*F*_ST_: 1.91% to 5.56%; *nS*_L_ in Africans: 1.12% to 7.08%; *nS*_L_ in Europeans: 2.25% to 5.39%; *nS*_L_ in East Asians: 2.25% to 5.93%). Furthermore, for 33/41 tissues examined, the H12 metric had the highest proportion of enhancers exhibiting evidence of recent selection. These results imply that human enhancers experienced selection at different time points in human history, with substantial evidence for soft selective sweeps in which multiple haplotypes increase in frequency within a population (Pennings and Hermisson 2006; Messer and Petrov 2013). However, we caution against overly interpreting differences in the proportions of enhancers across metrics as differences in the action of selection across different time periods since they likely vary in power to detect selection.

**Figure 2.**
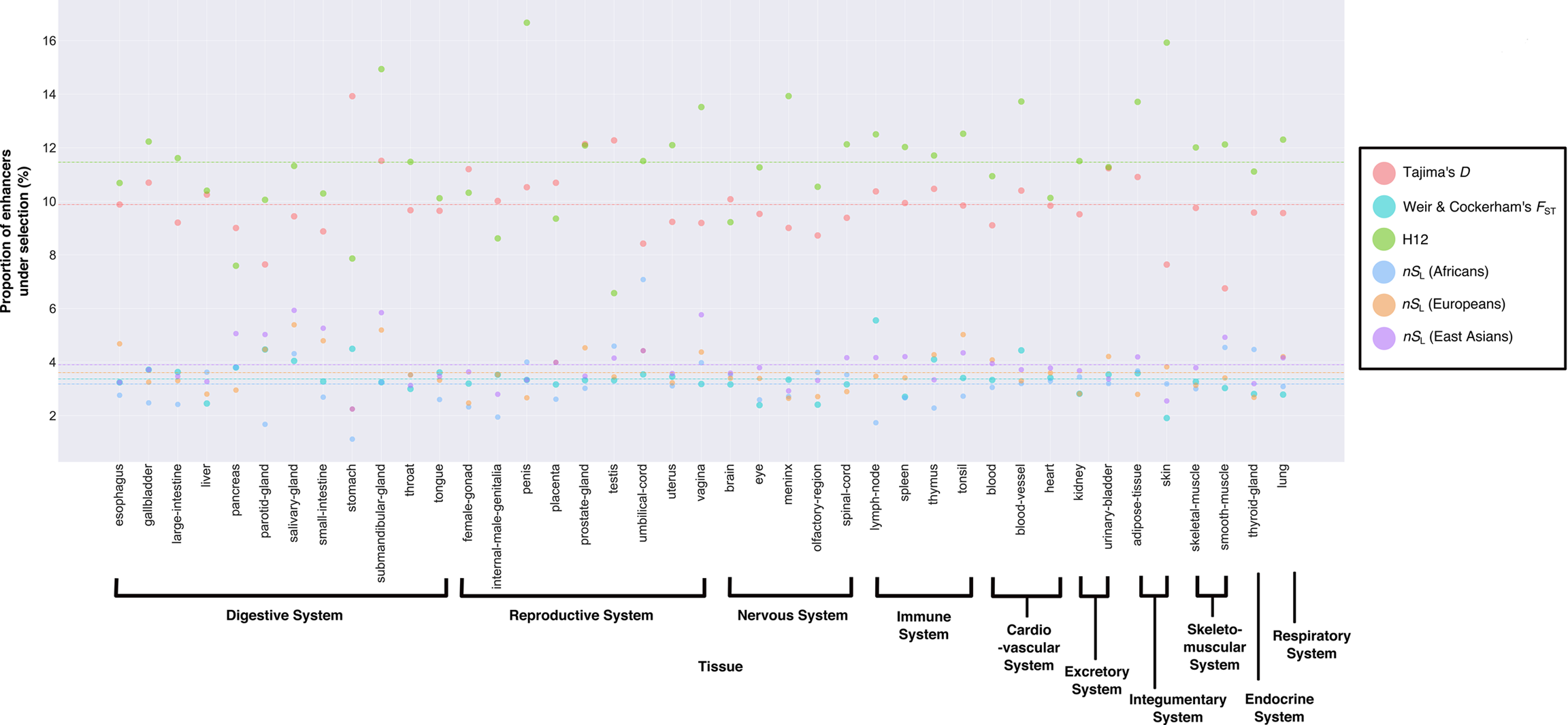
Proportions of enhancers exhibiting significant deviations from neutral expectations for different recent positive selection metrics across 41 tissues. Each dot represents the proportion of enhancers (Y-axis) that exhibited significant deviation from the neutral model (i.e., *p*-value < 0.05) in a given tissue (X-axis). Differently colored dots correspond to the different selection metrics used (Salmon: Tajima’s *D*; Teal: *F*_ST_; Green: H12; Blue Gray: *nS*_L_ (Africans); Orange: *nS*_L_ (Europeans); Purple: *nS*_L_ (East Asians)). For each metric, the *p*-value for the observed value for an enhancer (i.e., the likelihood under neutral expectations of obtaining a value as or more extreme as the observed value) was assessed by comparing to 10,000 simulated values calculated on sequences generated from the neutral simulations. Differently colored horizontal lines correspond to the average of all proportions calculated for all 41 tissues according to the different selection metrics used (Salmon: Tajima’s *D*; Teal: *F*_ST_; Green: H12; Blue Gray: *nS*_L_ (Africans); Orange: *nS*_L_ (Europeans); Purple: *nS*_L_ (East Asians)). Note the 1) differences in the proportion of enhancers exhibiting significant deviations from neutral expectations across tissues in any given metric, and 2) similarly, differences in the proportion of enhancers with significant deviations from the neutral model across all metrics in any given tissue.

### Enhancers that exhibit evidence of recent positive selection putatively regulate the activities of genes with immunity-related functions

To study the functions of the enhancers that show evidence of recent positive selection, we carried out enrichment analysis on the putative target genes associated with them (these results can be found in the Figshare repository). We found that there was little functional enrichment among the enhancers identified by *F*_ST_, Tajima’s *D* and H12 (Table 1). However, many tissues had multiple significant enriched GO terms among the *nS*_L_ hits (Table 1). For the *F*_ST_, Tajima’s *D* and H12 metrics, we also collapsed all enhancers irrespective of their tissues and carried out the same analyses: we found similar patterns as described above, with no significant GO terms for the *F*_ST_ metric and very few GO terms for Tajima’s *D* and H12 metrics (these results can be found in the Figshare repository). The majority of top 10 most frequent significant GO terms among all tissues for the *nS*_L_ metric were immunity-related (Table 2–4). In addition, we also carried out the same analyses on the putative target genes of enhancers with no evidence of recent positive selection according to the *nS*_L_ metric and found that none of the tissues possessed any significantly enriched GO terms: the only exception was a single Cellular Component term in the parotid-gland for *nS*_L_ metric in Europeans (GO:0044444: cytoplasmic part, adjusted *p*-value = 0.047). These results suggest that approximately 20,000–10,000 years ago, enhancers that putatively regulate the activities of immunity-related genes in multiple tissues underwent selection, a finding consistent with the hypothesis that human populations faced novel local selective pressures as they moved into new environments (Balaresque et al. 2007; Fumagalli et al. 2011).

**Table 1.**
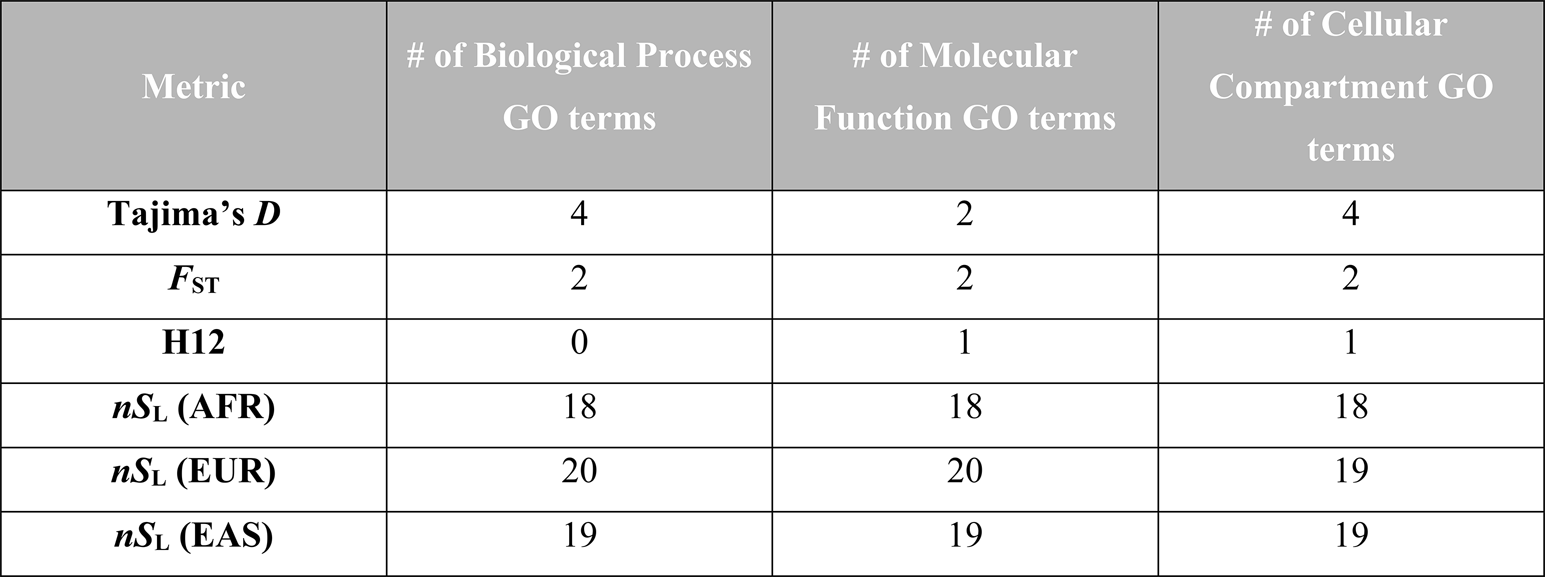
Number of tissues with one or more significantly enriched GO Terms.

**Table 2.**
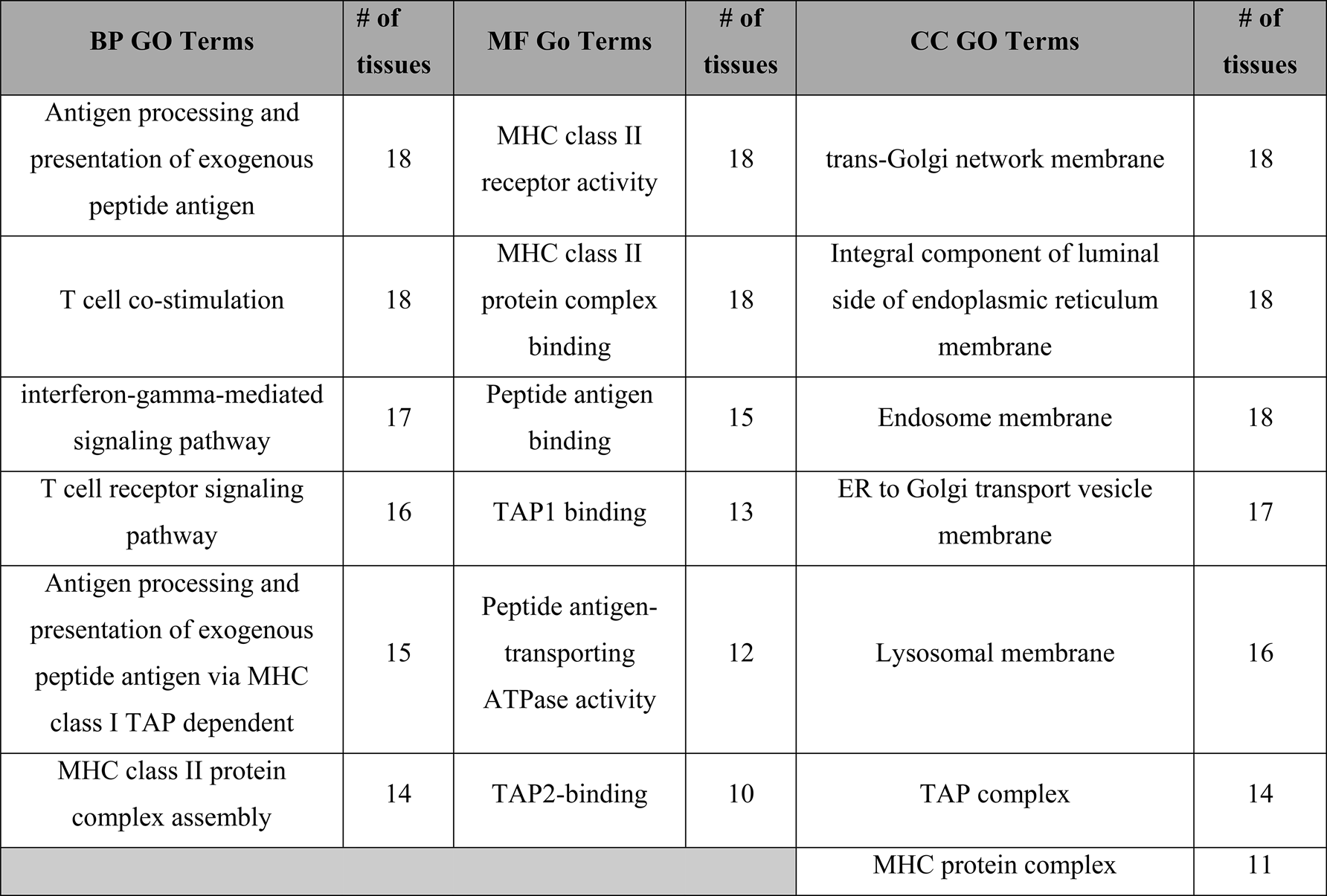
Most frequently-occurring GO terms among enhancers with evidence of recent positive selection by *nS*_L_ across tissues in Africans.

**Table 3.**
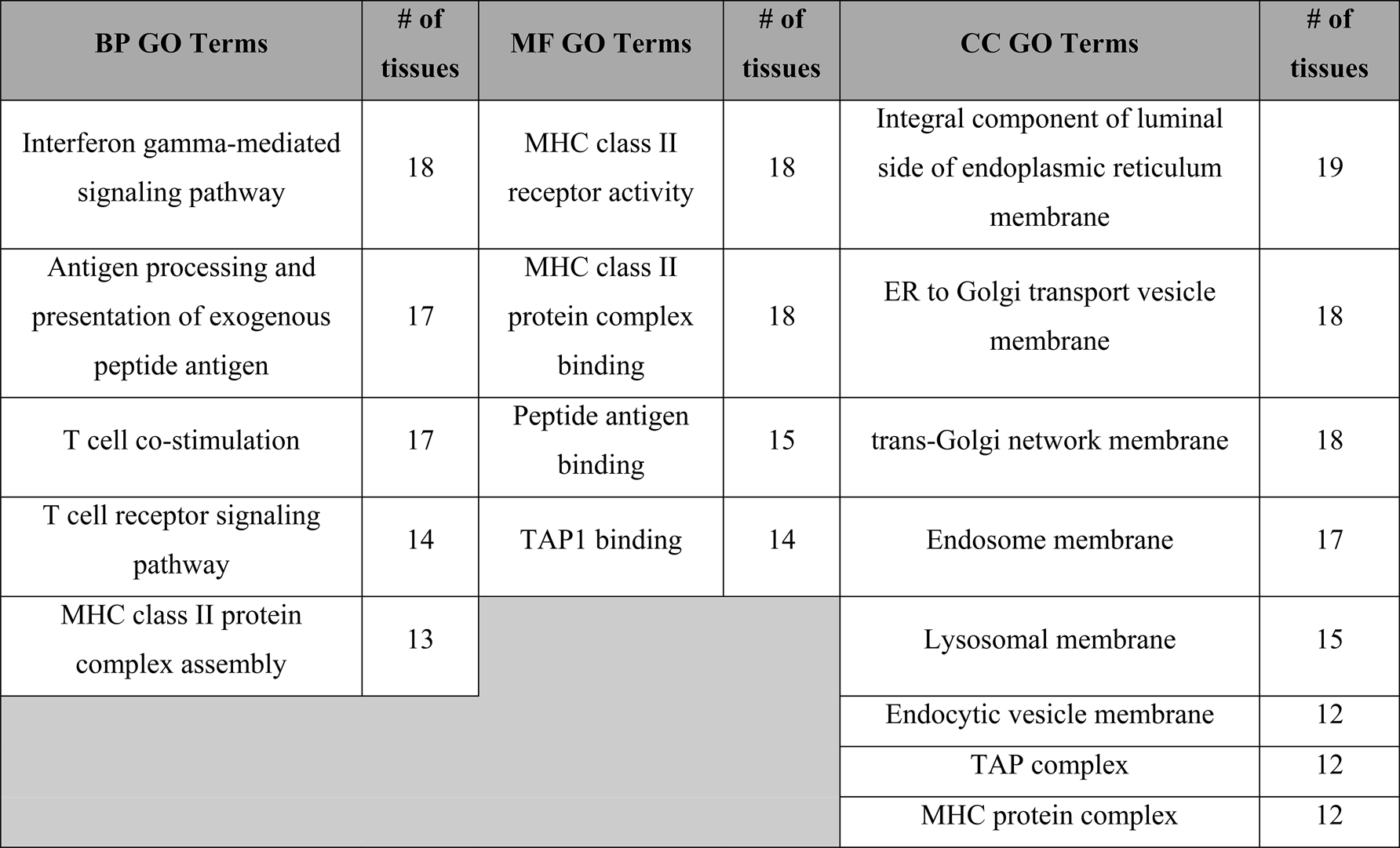
Most frequently-occurring GO terms among enhancers with evidence of recent positive selection by *nS*_L_ across tissues in Europeans.

**Table 4.**
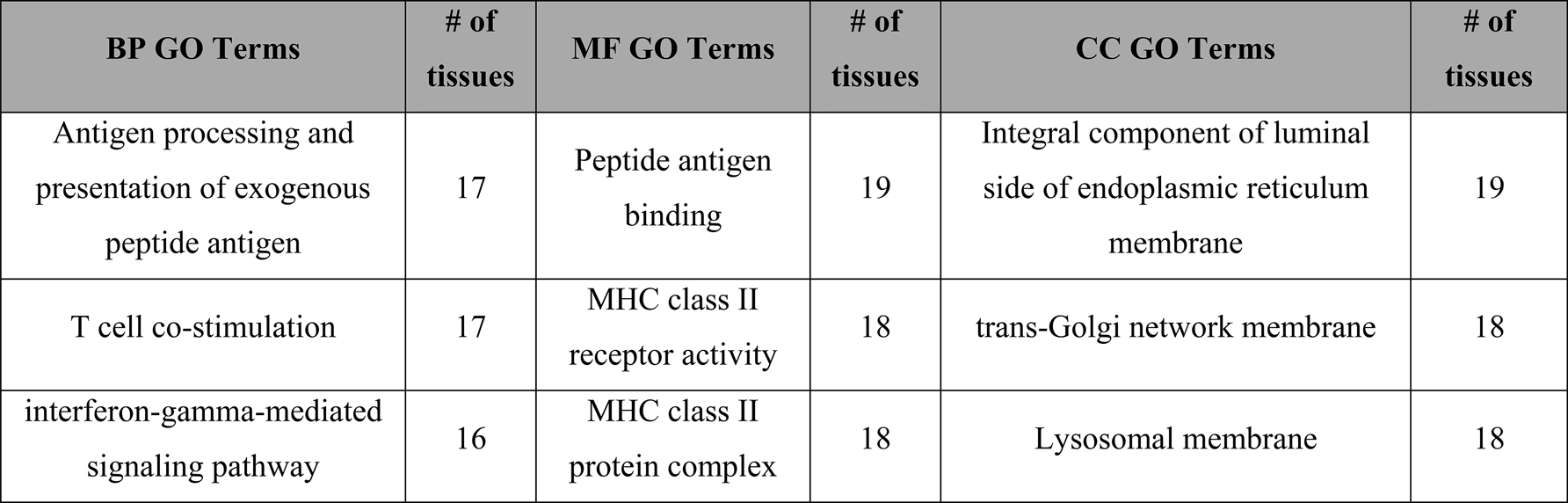
Most frequently-occurring GO terms among enhancers with evidence of recent positive selection by *nS*_L_ across tissues in East Asians.

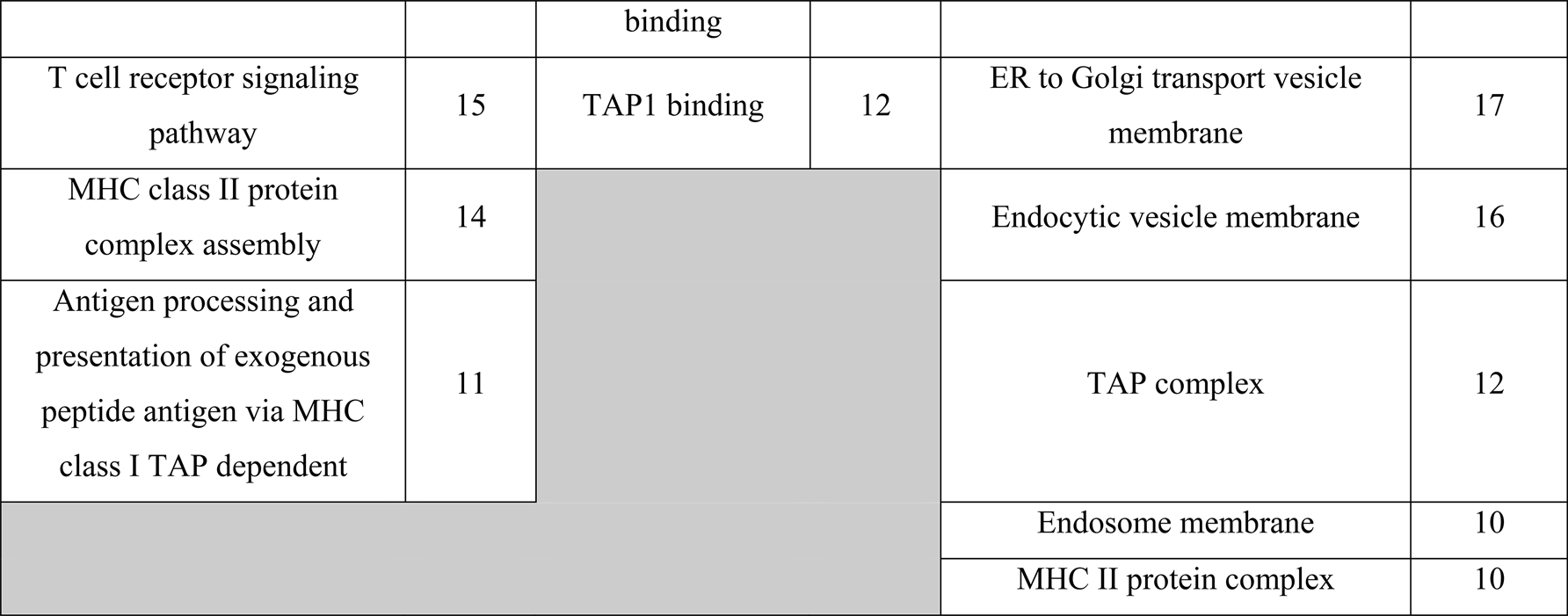

We also quantified the semantic similarity (SS) of the functionally enriched terms among the tissues to determine if the pattern of enrichment for immunity-related functions was shared across tissues (Figure S2a-i). The pairwise SS values calculated using the *TopGO* GO terms associated with the *nS*_L_ metrics were consistently very high (BP: Africans: 0.687 to 1.000; Europeans: 0.786 to 1.000; MF: Africans: 0.779 to 1.000; Europeans: 0.715 to 1.000; East Asians: 0.902 to 1.000; CC: Africans: 0.873 to 1.000; Europeans: 0.864 to 1.000; East Asians: 0.853 to 1.000): one exception to this general trend were the SS scores calculated on the Biological Process terms in East Asians (from 0.208 to 1.000) (Figure S2c). To determine if the range of SS values we observed were unusually high, we compared each pairwise SS value to the 95^th^ percentile value of the SS values calculated on the 1,000 randomly sampled GO terms for the same pair of tissues (these values, as well as the empirical SS values, can be found in the Figshare repository). We found that most of the empirical SS values were higher than the 95^th^ percentile values of the SS values calculated on the randomly sampled GO terms, with the exception of pairwise comparisons involving liver for the Biological Process terms in East Asians (Figure S2c). In other words, most pairs of tissues were associated with semantically similar GO terms, suggesting that patterns of functional enrichment were very similar across tissues. The sole exception to this were 13 comparisons involving the liver, which had three additional biological terms in East Asians (GO:0071294: cellular response to zinc ion, adjusted *p*-value = 0.005; GO:0046597: negative regulation of viral entry into host cell, adjusted *p*-value = 0.005; GO:007126: cellular response to cadmium ion, adjusted *p*-value = 0.014), which were found only in the liver. Overall, our analyses suggest that the enrichment for immunity-related functions found in putative target genes of enhancers with evidence of recent positive selection is not confined to specific tissues, but represents a general trend.

### Enhancers that exhibit evidence of recent positive selection are not enriched for transposable element origins

To examine whether enhancers that exhibit statistically significant signatures of recent positive selection tend to have arisen from TEs, we compared the proportions of TE overlap between enhancers that exhibit evidence of recent positive selection versus enhancers with no evidence of recent positive selection across all 41 tissues. Of the 41 tissues examined, only the lung (χ^2^ = 12.859, adjusted *p*-value = 0.002) and the female gonad (χ^2^ = 8.632, adjusted *p*-value = 0.020) displayed statistically significant differences in terms of the proportions of TE-overlapping regions between the two groups of enhancers for the H12 metric (Figure S3a-f). Moreover, in these two tissues, enhancers with evidence of recent positive selection showed lower proportions of TE overlap than enhancers with no such evidence. Overall, these results suggest that enhancers that have arisen from TEs were not preferential targets of recent positive selection.

### Enhancers active in testis and brain show different patterns of recent evolution

To test the hypothesis that signatures of selection during recent human history differed across enhancers active in different tissues, we compared the distributions of the four metrics across all 820 pairs of the 41 tissues (these results can be found in the Figshare repository). The number of tissue pairs exhibiting different distributions for Tajima’s *D* was 22 / 820, for *F*_ST_ was 5 / 820, for H12 was 56 / 820, and for *nS*_L_ was 2 / 820 in Africans, and 0 / 820 in Europeans and East Asians (Figure 3a-f). All tissue pairs exhibiting significantly different distributions involved either brain or testis, with the exception of two *nS*_L_ comparisons (Africans: Adipose-tissue and Large-intestine: *p*-value = 0.014; Adipose-tissue and Thymus: *p*-value = 0.007). Specifically, brain enhancers exhibited significantly different distributions for Tajima’s *D*, *F*_ST_, and H12 compared to enhancers from 5, 2, and 23 other tissues (Figure 3a-c); similarly, testis enhancers exhibited significantly different distributions for Tajima’s *D*, *F*_ST_ and H12 compared to enhancers from fifteen, three, and thirty other tissues (Figure 3a-c). We note that the high fractions of enhancers under selection in brain and testis are unlikely to be solely due to the high number of enhancers found in these tissues; specifically, there are several other tissues that have high number of enhancers (Table S1) and yet fail to exhibit statistically significant pairwise comparisons with other tissues.

**Figure 3.**
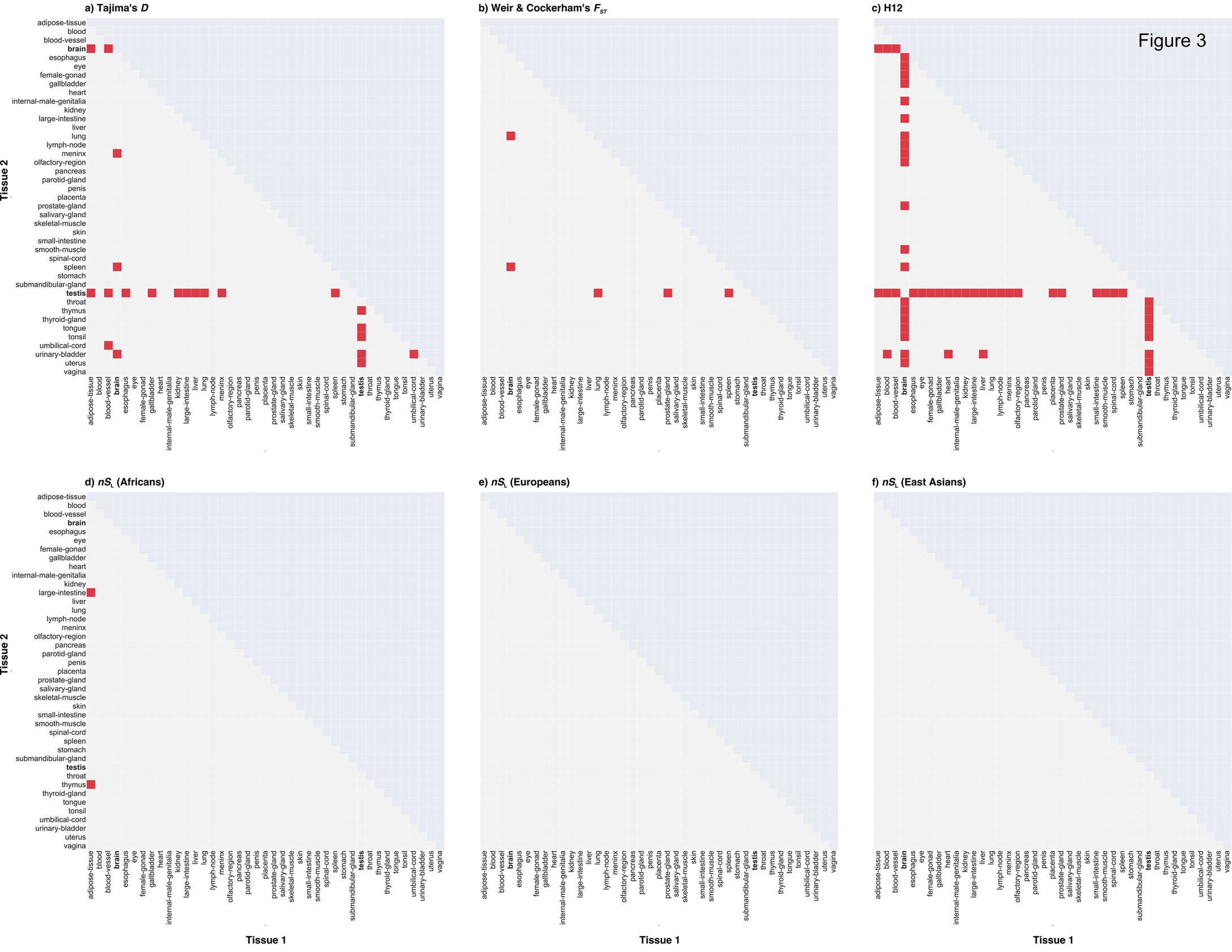
Pairwise comparisons of the patterns of recent evolution among enhancers from 41 tissues. Each graph shows the results of the pairwise Kolmogorov-Smirnov tests carried out between all pairs of 41 tissues (shown on the X- and Y-axes) for all recent positive selection metrics. Each cell represents a pairwise comparison between two specific tissues: red-filled cells represent pairwise comparisons that are statistically significant (i.e., adjusted *p*-value < 0.05) and empty cells represent non-significant pairwise comparisons.

In general, the significant differences in the distributions of these metrics between testis or brain and other tissues were due to differences in the magnitude of the peaks of the distributions and/or differences in shapes of the distributions rather than from shifts in the range of the distributions. For all metrics, most pairs of tissues with significant differences had very subtle shifts in the distributions, and in cases where the shifts were more noticeable, brain and testis enhancers’ distributions tended to be shifted to the left (for *F*_ST_; Figure S5) or the right (for Tajima’s *D*; Figure S4), both in the direction suggestive of lower levels of positive selection. The nature of the differences in the peaks of the magnitude varied for each metric: for *F*_ST_, the peaks for both brain and testis were consistently higher compared to those of the other tissues. In contrast, for Tajima’s *D*, the peaks for brain and testis were consistently lower than the other tissues being compared; the sole exception was the comparison between brain and meninx and urinary-bladder, in which the peaks were almost equal or slightly higher for the brain, respectively. In addition, for Tajima’s *D*, there were differences in the shapes of the distribution (Figure S4). For H12, differences in the shape of the distributions also contributed to the statistical differences we observed between brain or testis and other tissues (Figure S6-7). These results imply that enhancers active in the brain and testis experienced different selective pressures during recent human history compared to enhancers active in other tissues.

We also ranked the tissues according to the median values of each metric (Figure S8) and the proportion of enhancers that exhibit statistically significant evidence of recent positive selection for each metric (Figure S9). Overall, we did not find any general patterns regarding the recent evolution of brain and testis, in terms of either the median values of the metrics of recent selection (Figure S8) or the proportion of enhancers that have putatively been under selection (Figure S9). Exceptions to the general trend were that testis and brain ranked 1st and 2nd, respectively, for median values of the H12 metric (Figure S8c) and that testis ranked 2nd in the proportion of enhancers with evidence of recent selection according to Tajima’s *D* and *nS*_L_ in Africans (Figure S9a & d).

### Evolution of tissue-specific enhancers differs from that of tissue-broad enhancers in the brain and testis

We next compared the patterns of recent evolution between tissue-specific and tissue-broad enhancers for each tissue. We found that the number of tissues with statistically significant differences in the distributions of the metrics between enhancers active in a single tissue and those active in multiple tissues was as follows, sorted from highest to lowest: 20/41 for H12, 8/41 for Tajima’s *D*, 8/41 for *F*_ST_ and 1/41 for *nS*_L_ in Africans and East Asians (Figure 4a & S10-15). Tissue-specific and tissue-broad enhancers in the brain and testis show statistically significant differences in the patterns of recent evolution for all metrics except *nS*_L_ (Tajima’s *D*: brain: 1.882e-07; testis: 9.523e-11; *F*_ST_: brain: 6.074e-07; testis: 1.639e-07; H12: brain: < 2.2e-16; testis: < 2.2e-16 (Figure 4b-d). Regarding the nature of the differences between the two groups of enhancers in these tissues, we found contrasting patterns for H12 compared to Tajima’s *D* and *F*_ST_: for Tajima’s *D* and *F*_ST_, we found that the interquartile values of the tissue-specific enhancers were lower (or higher for Tajima’s *D*) than those of the tissue-broad enhancers in brain and testis, whereas those values were consistently higher for the tissue-specific enhancers compared to the tissue-broad enhancers in the same tissues for the H12 metric. These results suggest that, in general, enhancers with different breadth of tissue activity did not differ in their patterns of recent evolution. Nevertheless, enhancers active only in the brain do show significant differences compared to those active in multiple tissues (including brain) and the same is true for testis, raising the possibility that the recent evolution of brain- and testis-specific enhancers may have been different from that of other tissue-specific enhancers.

**Figure 4.**
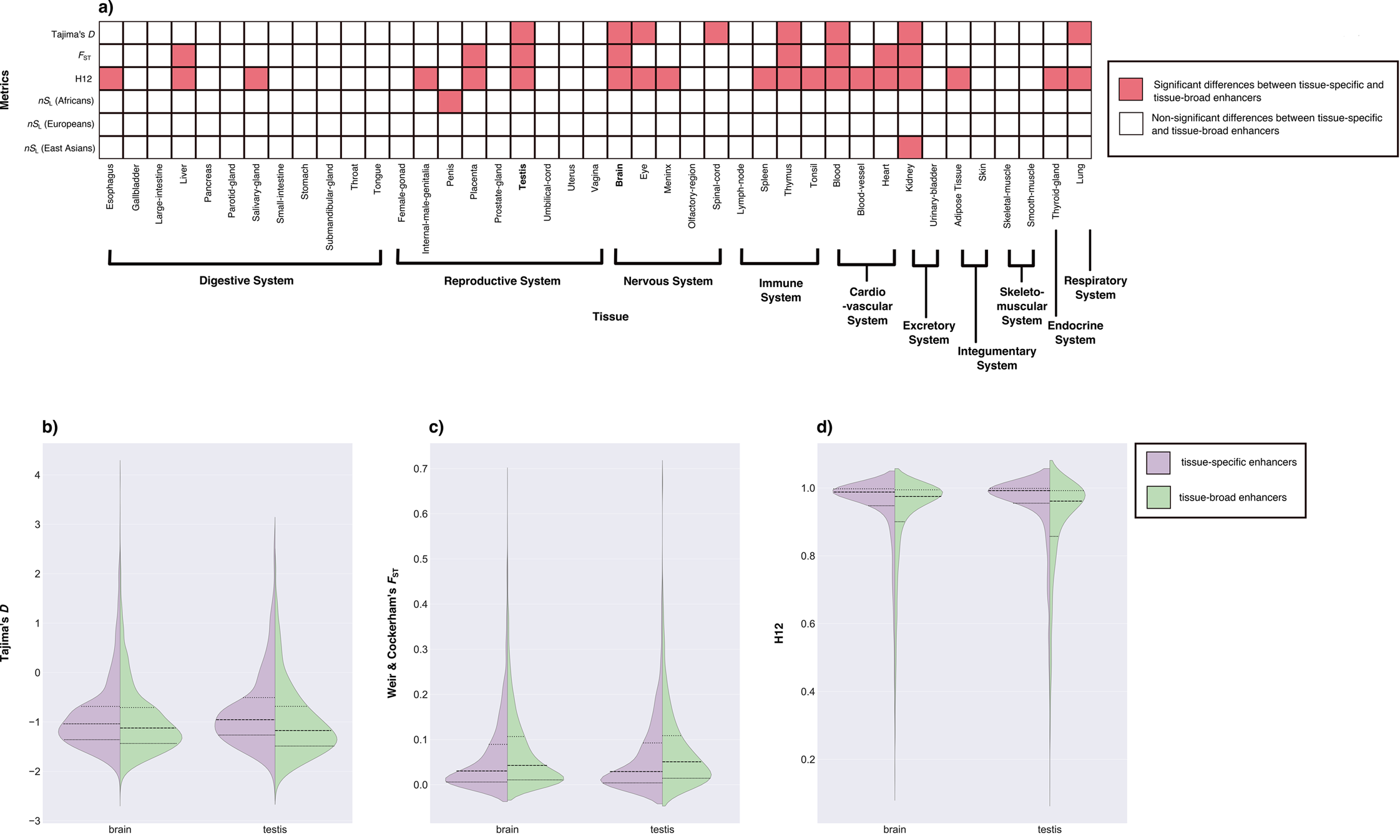
Comparisons of patterns of recent evolution between tissue-specific and tissue-broad enhancers. (a) The grid panel depicts which tissues exhibit significantly different distributions of metrics between tissue-specific and tissue-broad enhancers. Each cell represents a comparison between tissue-specific and tissue-broad enhancers in a given tissue: pink-filled cells represent the comparisons that are statistically significant (i.e., adjusted *p*-value < 0.05) and empty cells represent non-significant comparisons. (b–d) Violin plots depicting the distributions of the metrics for tissue-specific and tissue-broad enhancers for brain and testis for (b) Tajima’s *D*, (c) Weir & Cockerham’s *F*_ST_, and (d) H12. In all cases, the distributions of tissue-specific and tissue-broad enhancers were significantly different (see also panel a).

### The recent evolution of testis enhancers associated with tissue-enriched genes differs from the evolution of enhancers associated with non-tissue-enriched genes

To determine whether there are differences in the patterns of recent evolution between enhancers associated with tissue-enriched genes versus those associated with non-tissue-enriched genes, we compared the distributions of the four metrics between brain and testis enhancers stratified by the breadth of expression of their target genes (these results can be found in the Figshare repository). There were significant differences for H12 (adjusted *p*-value =3.667e-06) and Tajima’s *D* (adjusted *p*-value = 0.004) in the testis (Figure 5a-b). More specifically, we found that the distribution of Tajima’s *D* values of the enhancers associated with testis-enriched genes were shifted to the left (i.e., more negative values) compared to the enhancers associated with non-testis-enriched genes (Figure 5a). The difference between the same groups of enhancers for the H12 metric was comparatively subtle, with the main difference being the magnitude of the peak (Figure 5b); the peak of the distribution of the enhancers associated with the non-testis-enriched genes was higher than that of the enhancers associated with testis-enriched genes. In contrast, there were no significant differences in the distributions of *F*_ST_ and *nS*_L_ values between these two categories of enhancers for either tissue and the distributions of H12 and Tajima’s *D* between the same categories of enhancers for the brain (Figure 5). These results imply that enhancers associated with genes with enriched expression in the testis have experienced different degrees of selection in the form of both hard and soft selective sweeps compared to enhancers associated with genes that do not show enriched expression in the same tissue.

**Figure 5.**
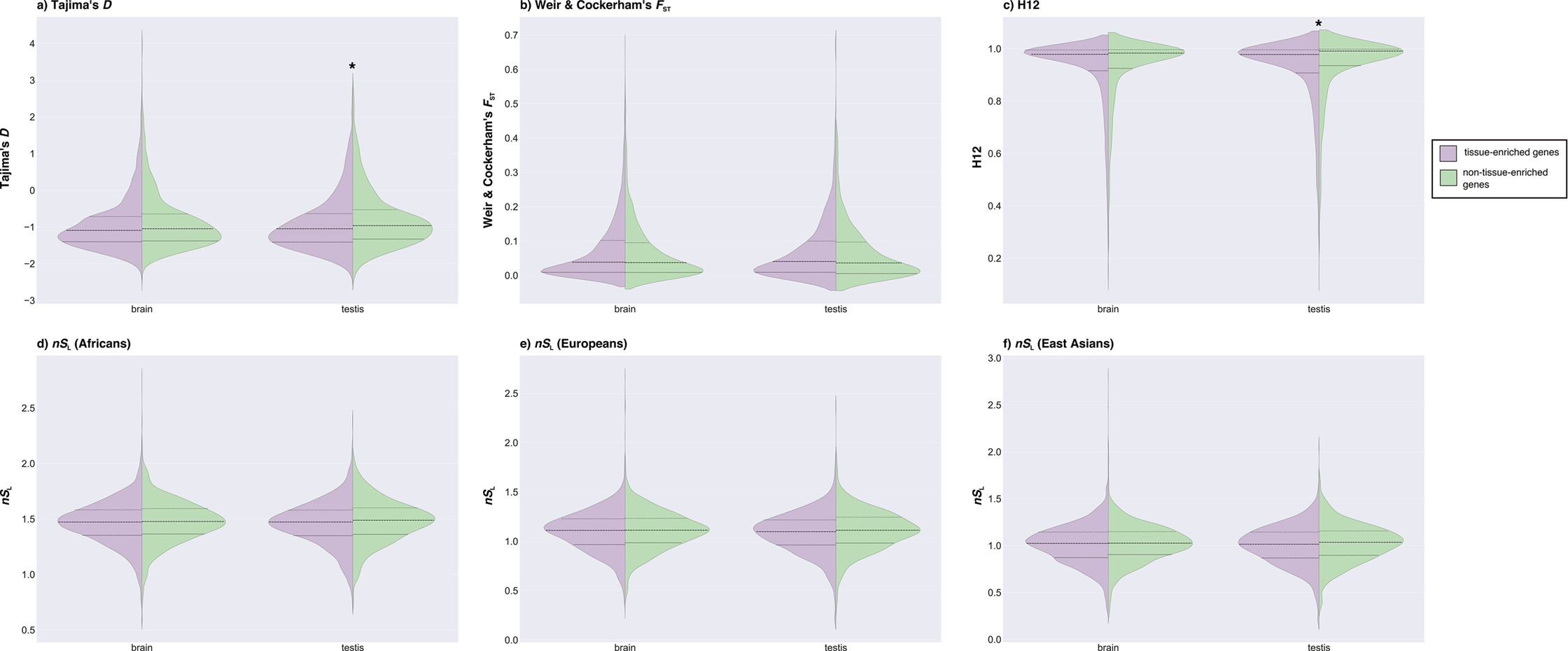
Comparisons of patterns of recent evolution between enhancers associated with tissue-enriched genes and enhancers associated with non-tissue-enriched genes in brain and testis. The violin plots depicting the distributions of the metrics for enhancers associated with tissue-enriched and non-tissue-enriched genes for brain and testis for: (a) Tajima’s *D* (b) Weir & Cockerham’s *F*_ST_ (c) H12 (d) *nS*_L_ (Africans) (e) *nS*_L_ (Europeans) (f) *nS*_L_ (East Asians). Plots with asterisks above them indicate significant pairwise comparisons.

### Brain and testis enhancers under recent positive selection are not significantly enriched for variants associated with complex human traits and diseases

To examine if genetic variants in brain and testis enhancers showing signatures of recent selection are associated with particular complex human traits and diseases, we queried the NHGRI-EBI GWAS catalog (MacArthur et al. 2017). We found that for all metrics, most or all enhancers did not harbor SNPs that are associated with any human traits (brain: Tajima’s *D*: 372/421; *F*_ST_: 131/150; H12: 430/437; *nS*_L_(Africans): 150/167; *nS*_L_ (Europeans): 144/161; *nS*_L_ (East Asians): 152/170; testis: Tajima’s *D*: 143/163; *F*_ST_: 47/52; H12: 102/103; *nS*_L_ (Africans): 64/72; *nS*_L_ (Europeans): 48/54; *nS*_L_ (East Asians): 60/65). In the few instances in which SNPs within enhancers showing evidence of selection were associated with human traits, these associations were with traits such as Alzheimer’s disease (brain: Tajima’s *D*), obesity-related traits (brain: Tajima’s *D*), and type 2 diabetes (brain: *nS*_L_ (Africans)) (Tables 5 & 6; the full list of overlapping GWAS hits, including variants in LD with those inside enhancers, can be found in the Figshare repository). We next compared the proportions of enhancers that overlapped with GWAS hits between enhancers that exhibit evidence of recent positive selection and enhancers that do not and did not find any significant difference between them. More specifically, we saw no statistically significant differences in the proportions of overlap with GWAS hits between enhancers with or without evidence of recent positive selection: the only exception was the H12 metric in the brain (χ^2^= 30.552, adjusted *p*-value = 1.950e-07), which showed depletion of GWAS hits among enhancers with evidence of recent positive selection (These results can be found in the Figshare repository). These results show that overall, enhancer SNPs under recent positive selection are not preferentially associated with specific human traits or complex human diseases.

**Table 5.**
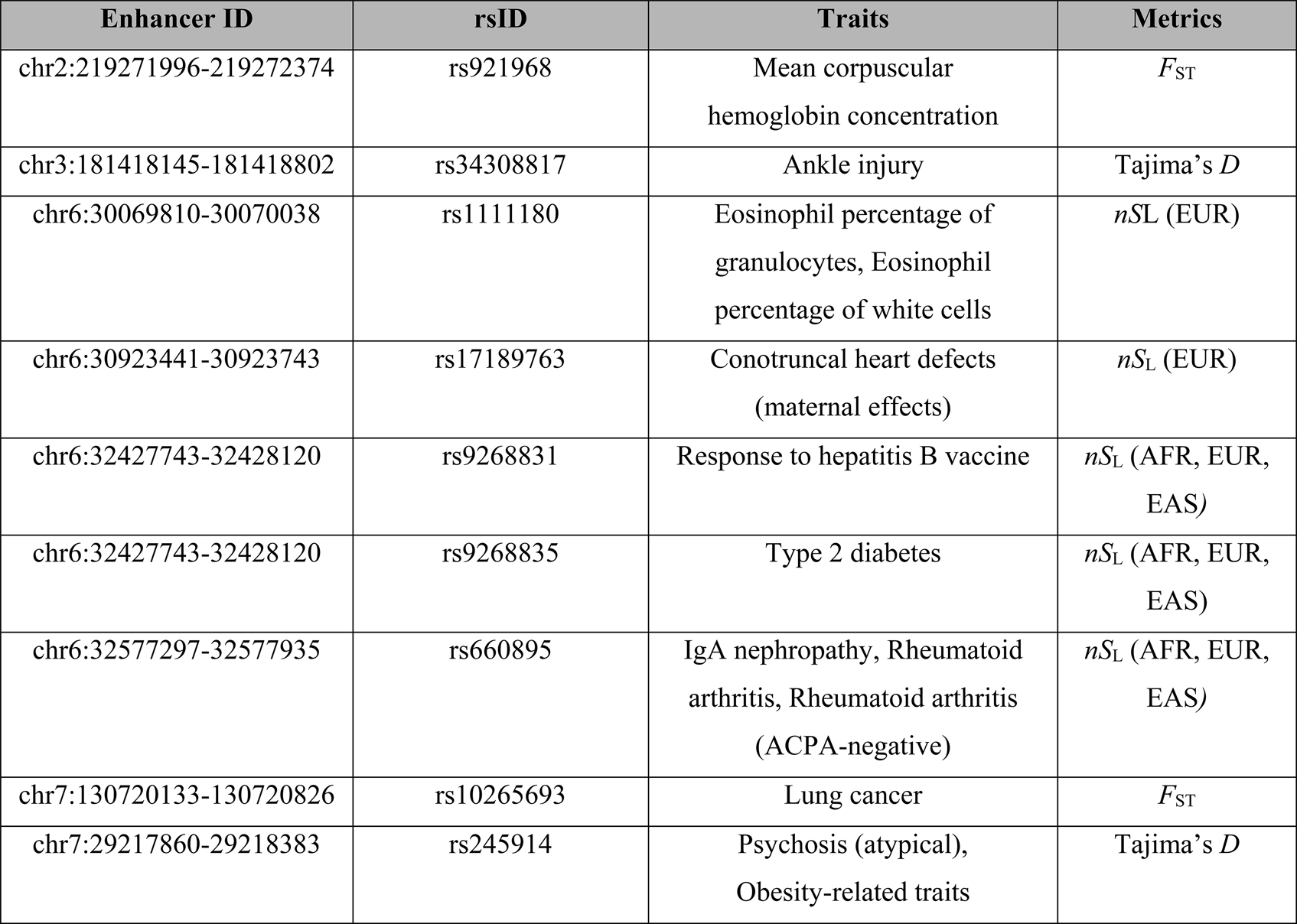
Complex human traits and diseases associated with variants within the brain enhancers that exhibit evidence of recent selection.

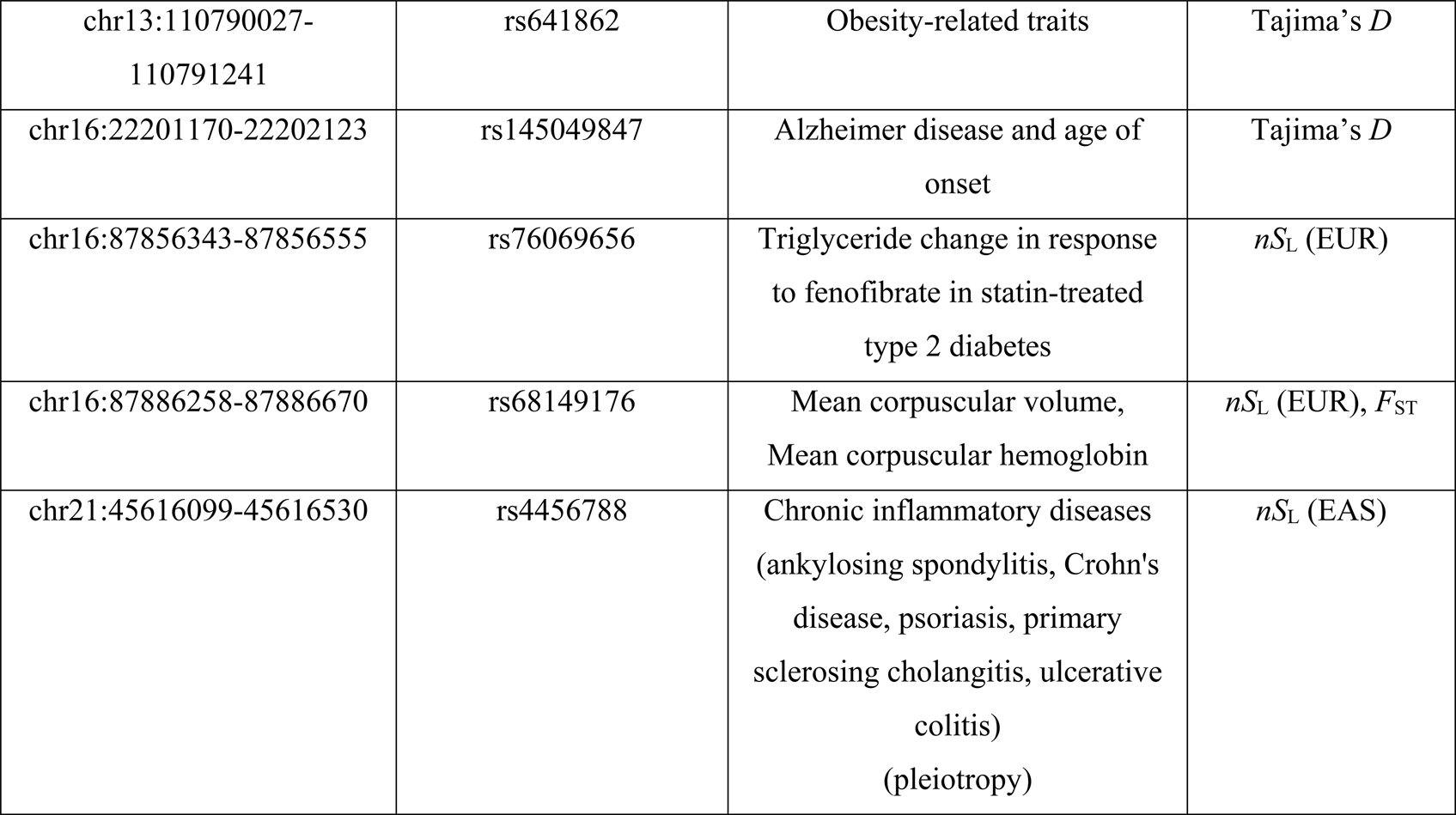

**Table 6.**
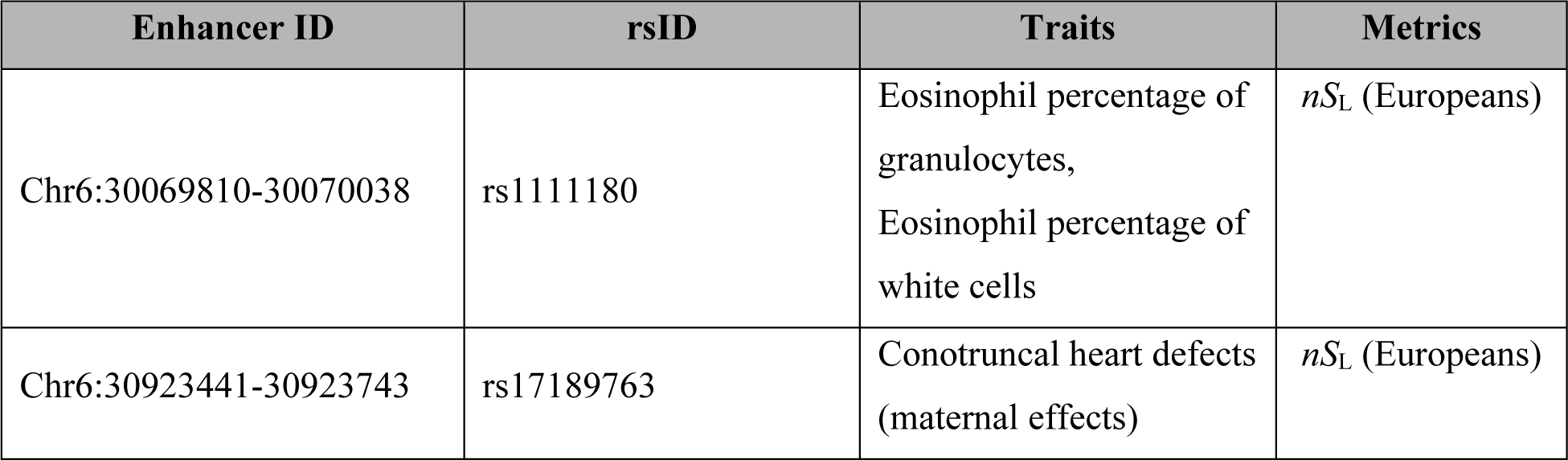
Complex human traits and diseases associated with variants within the testis enhancers that exhibit evidence of recent selection.

## Discussion

In this study, we calculated four different metrics that detect distinct genomic signatures of recent selection on the enhancers active in 41 human tissues and compared the empirical values of these metrics to those calculated on neutrally simulated sequences. We found that across all tissues and metrics, approximately 5.90% of enhancers exhibit significant evidence of recent positive selection. We also found that the putative target genes of such enhancers are enriched for immunity-related functions, and that this enrichment pattern is shared across multiple tissues. Furthermore, enhancers active in the brain and testis exhibited significantly different patterns of recent evolution compared to enhancers in other tissues.

Across tissues, we observed variation in the proportions of enhancers that exhibited signatures of selection across the different metrics (Figure 2). Since the origin of modern humans approximately 250,000 - 200,000 years ago, there have been multiple distinctive events in human history, including the out-of-Africa migration approximately 75,000 - 50,000 years ago, the Eurasian split that occurred approximately 45,000 - 36,000 years ago, and the Agricultural Revolution approximately 20,000 - 10,000 years ago, all of which would have likely exposed the human populations to novel selective pressures (e.g., pathogens, dietary changes) (Sabeti et al. 2006; Karlsson et al. 2014). Adaptations in response to such changes would have likely altered the allele frequencies in both the genic and regulatory regions. Furthermore, it has been previously shown that different metrics of selection are sensitive to distinct types of genomic signatures (Sabeti et al. 2006; Voight et al. 2006). For example, Tajima’s *D* is most suitable for detecting events that occurred approximately 250,000 - 200,000 years ago, *F*_ST_ is most suitable for identifying selection events that occurred as humans left Africa and were exposed to novel environments around the world (i.e., 75,000 - 50,000 years ago), whereas *nS*_L_ is most suitable for detecting selection events that happened approximately 20,000 - 10,000 years ago (Sabeti et al. 2006; Voight et al. 2006). Therefore, one possible interpretation is that within any given tissue, the differences in percentages of enhancers under selection between metrics reflect temporal variation in the action of selection during human history.

While the variation in the proportion of enhancers with significant evidence of recent selection across different metrics could represent temporal variation in the action of selection, it is also likely that these metrics have different power to detect selection. For instance, Tajima’s *D*, a metric that detects excess of rare alleles in a region of interest, has been shown to be most sensitive to selective sweeps that have resulted in almost complete fixation (i.e., allele frequencies = 100%) of the target locus and conversely has limited power to detect actions of selection that have resulted in incomplete fixation of the allele (Sabeti et al. 2006; Ferrer-Admetlla et al. 2014). In contrast, *nS*_L_, a haplotype-based metric, best detects ongoing or incomplete hard selective sweeps resulting in intermediate allele frequencies (i.e., allele frequencies = 60 - 80%) and rapidly loses power as allele frequencies increase to 100% (Ferrer-Admetlla et al. 2014). Thus, the observed differences in the proportions of enhancers with evidence of recent positive selection across the metrics also reflect the disparity in power of the metrics to successfully detect incidences of selection that happened at a particular time period. Consequently, the proportions of enhancers with evidence of recent selection across metrics cannot be directly compared to each other. Distinguishing whether our results are best explained by temporal variation in selection or by differences in the power of our metrics could be achieved via simulations in which selective events are introduced at specific time points and result in a fixed proportion of regions being selected. This is an important future research direction that has potential to shed light into the tempo of selection in the course of recent human evolution.

An additional result of our study is that the putative target genes of enhancers exhibiting significant evidence of recent positive selection according to the *nS*_L_ metric are enriched for immunity-related functions (Tables 2–4). Haplotype-based metrics such as *nS*_L_ are known to be sensitive to signatures resulting from selection that occurred approximately 10,000 - 20,000 years ago, which corresponds to the incidence of the Agricultural Revolution (Voight et al. 2006). The advent of farming practices, communal living in settlements, and migrations of farmers across the globe resulted in an increase of numbers and densities of humans in any given location, likely facilitating the spread of pathogens (Varki and Gagneux 2009; Page et al. 2016; Nielsen et al. 2017). In addition, recent studies suggest that selection on *cis*-regulatory regions, such as enhancers, might have been important in driving adaptation of modern human populations to distinct environments, due to their modular organization: change of expression pattern in one temporal or spatial context can often occur without affecting others, which could contribute to phenotypic changes without incurring negative pleiotropic effects (Carroll 2005; Wray 2007). Therefore, it is possible that around 10,000 - 20,000 years ago, enhancers regulating the activities of immunity-related genes underwent selection to allow refined fine-tuning of host defense processes in response to the stronger pressure from pathogens resulting from increased human population sizes.

Perhaps the most striking result of our study is that brain and testis enhancers exhibited different patterns of recent evolution compared to enhancers of other tissues (Figure 3 & S4-7). For both brain and testis, the high number of enhancers included in these tissues could partly explain our results. Brain, with 4,883 enhancers, has the most enhancers of all the tissues examined, while testis, with 1,621 enhancers, is ranked 7th. The effect of the large number of enhancers included in these tissues is reflected in the differences in the magnitudes of peaks of the distributions of the metrics between brain and testis and other tissues (Figure S4-7). However, we also observed other types of differences in comparisons of the distributions of these metrics between brain and testis and other tissues: while subtle, some metrics (Tajima’s *D*, *F*_ST_; Figure S4-5) showed shifts of the distributions of brain and testis enhancers towards weaker signatures of recent positive selection compared to other tissues, suggesting that the significant pairwise differences we see are not solely caused by disparities in the number of enhancers within tissues. In addition, there are several other tissues (e.g., lung, spleen, blood) whose numbers of enhancers are nearly on par with those of the brain and more than the testis; however, these tissues did not show significant differences in the distribution of these metrics compared to other tissues, further suggesting that the observed differences in selection are likely biological.

Why are patterns of selection different for brain and testis enhancers? Answering these questions is challenging without additional functional experiments that shed light on the phenotypic effects of the selected variants. In the case of the brain, existing evidence suggests that the brain size of modern humans has not changed substantially in the last 250,000 - 300,000 years (Neubauer et al. 2018). Similarly, expression patterns of genes expressed in neural tissues (e.g., brain, cerebellum) show low levels of divergence across both species and within humans, suggesting that the overall structure of the neural network is highly conserved (Khaitovich 2005; Brawand et al. 2011). In contrast, brain shape has gradually changed in modern humans, reaching the present-day variation human variation about 100,000 - 35,000 years ago, but whether this has anything to do with positive selection being relatively weaker in the brain compared to other tissues is pure speculation.

In the case of testis, our results suggest that enhancers active in the testis often show weaker signatures of positive selection compared to enhancers in other tissues. Previous studies have shown that testis exhibits the highest degree of expression pattern divergence between species, likely due to either strong action of positive selection on reproductive processes (Khaitovich 2005; Brawand et al. 2011), relaxation of purifying selection in the testis compared to other tissues (Gershoni and Pietrokovski 2014), or both. Unfortunately, our results are not directly comparable with studies measuring gene expression divergence between species due to the difference in the features being compared and the temporal window examined; previous studies have looked at the divergence of the overall transcriptome, which is influenced not just by changes in enhancer sequences but by other types of changes and other regulatory elements as well, whereas we have specifically looked at changes in allele frequency and haplotypes in enhancer regions. In addition, previous studies have looked at divergence of the expression patterns between species that diverged several millions of years ago, whereas we have examined selection events that have occurred in more recent evolutionary times, on the order of hundreds or tens of thousands of years. Therefore, it is possible that there has been temporal variation (i.e., ancient vs. recent) in the occurrence of selection events on these enhancer regions. It is worth noting that Khaitovich et al. (Khaitovich et al. 2005) also found that testis had the most significant reductions in diversity of expression (i.e., low inter-individual variation in expression patterns) compared to other tissues examined, which points to differences in the action of selection over various temporal windows within the testis. In short, differences between our results and those of previous studies can be explained by differences in actions of ancient versus recent selection.

## Data Availability

Additional data from this study are available through the Figshare data repository (10.6084/m9.figshare.7629434; note that this link will become available upon publication).

## Acknowledgments

This research was supported by the March of Dimes through the March of Dimes Prematurity Research Center Ohio Collaborative, the Burroughs Wellcome Fund, the Guggenheim Foundation, and the National Institutes of Health (R01GM115836, R35GM127087). This work was conducted, in part, using the Advanced Computing Center for Research and Education at Vanderbilt University.

