## Supplementary Information for "Signatures of recent positive selection in enhancers across 41 human tissues"

**Supplementary Material**

File S1: Extended Methods
Figures S1-15
Table S1: Number of autosomal enhancers included in the 41 tissues
Table S2: Proportions of enhancers with significant evidence of recent positive selection according to the calculated metrics in all 41 tissues

**Materials Available Through The Figshare Repository**

File_1: Average lengths of enhancers used in our neutral simulations for all 41 tissues
File_2: Recombination rate parameters used in our neutral simulations for all 41 tissues
File_3: Compressed directory containing raw data CSV files: for each enhancer within each tissue, lists the values of the calculated metrics and the *p*-values computed from the neutral simulations
File_4-20: *TopGO* enrichment analysis results for each metric

- File_4-6: Biological Process (BP) terms for Africans, Europeans, East Asians
- File_7-9: Molecular Functions (MF) terms for Africans, Europeans, East Asians
- File_10-12: Cellular Components (CC) terms for Africans, Europeans, East Asians
- File_13-14: BP terms for Tajima's $D$, $F_{ST}$
- File_15-17: MF terms for Tajima's $D$, $F_{ST}$, H12
- File_18-20: CC terms for Tajima's $D$, $F_{ST}$, H12

Table_1-4: *TopGO* enrichment analysis results carried out for Tajima's $D$, $F_{ST}$, H12 by collapsing the tissues
Table_5-13: Pairwise Semantic Similarity values calculated for the $nS_L$ metrics
Table_14-19: Chi-square test results for the TE enrichment analyses
Table_20-25: KS test results for pairwise comparisons of distributions of metrics between tissues
Table_26-31: KS test results for comparisons of distributions of metrics between tissue-specific and tissue-broad enhancers for each tissue

Table_32-33: KS test results for comparisons of distributions of metrics between enhancers associated with tissue-enriched and non-tissue-enriched genes for brain and testis
Table_34-35: GWAS hits overlapping with both variants within enhancers and variants in LD with those inside enhancers for brain and testis
Table_36-37: Chi-square test results for GWAS hits enrichment analyses for brain and testis

**File S1. Extended Methods**

*1000 Genomes Project Phase 3 Genetic variation Dataset*

The compressed genotype files (*.vcf.gz) and tab-delimited index (*.tbi) files for all autosomal chromosomes were downloaded from:

http://ftp.1000genomes.ebi.ac.uk/vol1/ftp/release/20130502/

We downloaded the human sample panel file that contains population information of each individual included in the genotype dataset from:

http://ftp.1000genomes.ebi.ac.uk/vol1/ftp/release/20130502/integrated_call_samples_v3.20130502.ALL.panel

The 2,504 individuals included in this dataset were divided into five major groups (i.e., 'super-populations') using information in the human sample panel file. We processed the above file to create three 'population files' for the three major populations of interest (Africans, Europeans, East Asians), in which each file contains the IDs of individuals belonging to a given population group. For subsequent analyses, we used genotype data of 1,668 individuals belonging to the three major populations of interest.

*FANTOM5 Enhancers Dataset*

We downloaded the BED files listing the entire set of FANTOM-predicted enhancers active in the 41 human tissues from:

http://enhancer.binf.ku.dk/presets/

All coordinates were converted from 0-based to 1-based before running the subsequent analyses. We also downloaded the Transcription Start Site (TSS)-enhancer associations dataset:

http://enhancer.binf.ku.dk/presets/enhancer_tss_associations.bed

*HapMap Phase II Genetic Map Data*

We downloaded the HapMap Phase II genetic map that had been lifted from build 35 to GRCh37 from:

ftp://ftp-trace.ncbi.nih.gov/1000genomes/ftp/technical/working/20110106_recombination_hotspots/HapmapII_GRCh37_RecombinationHotspots.tar.gz

We carried out a linear interpolation to annotate the 1000 Genomes Project Phase III variants that are not included in the HapMap Phase II project with the estimated recombination rates (cM/Mb).

*Calculation of Average Recombination Rates per Tissue*

To calculate the average recombination rate for each tissue, we took all the autosomal enhancers in a given tissue and for each enhancer, annotated the variants with the recombination rates for that site using the data from the preceding section ("HapMap Phase II Genetic Map Data") and calculated the average recombination rate (cM/Mb) for the enhancer. We next took the average of the average values of recombination rates for all the enhancers in a given tissue to represent the recombination rate (cM/Mb) for that tissue. Finally, we further converted the average recombination rate per tissue to parameters *SLiM* can recognize (i.e., crossover per bp) by multiplying $10^{-8}$.

*VCFtools* (v.0.1.13)

*VCFtools* was used to calculate Weir & Cockerham's $F_{ST}$ and to create new VCF files containing genotype data within each enhancer region in the FANTOM dataset. To calculate global $F_{ST}$ for each enhancer, we used the following command:

**./vcftools --gzvcf input.vcf.gz-- chr # --from-bp start_pos --to-bp end_pos --remove-indels --weir-fst-pop AFR_samples_list --weir-fst-pop EUR_samples_list --weir-fst-pop EAS_samples_list --out output**

'input.vcf.gz' file refers to the compressed genotype files, sorted by chromosomes, downloaded from the 1000 Genomes Project ftp site. '#' indicates the chromosome id. 'start_pos' and 'end_pos' refer to the start and end positions of the enhancer, in base pairs. We used the weighted $F_{ST}$ values recorded in the *.log file created from the above command. Next, we created new VCF files that contain the genotype data for each enhancer using the coordinates recorded in the FANTOM BED files:

**./vcf --gzvcf input.vcf.gz --chr # --from-bp start_pos --to-bp end_pos --remove-indels --recode --recode-INFO-all --out output**

This command creates a new VCF (output.recode.vcf) file with genotype data encompassing the regions defined by the '--from-bp' and '--to-bp' arguments. Finally, we created separate VCF files that contain the genotype data for the region that spans 50kb up and downstream of a given enhancer for each population as follows:

**./vcf --gzvcf input.vcf.gz --chr # --from-bp start_pos_50kb --to-bp end_pos_50kb --remove-indels --min-alleles 2 --max-alleles 2 --keep pop_list --recode --recode-INFO-all --out output**

'start_pos_50kb' and 'end_pos_50kb' refer to positions 50kb upstream and downstream of the start and end positions of an enhancer as recorded by FANTOM, respectively. 'pop_list' is the file that lists all the individuals belonging to a given population group. The '--min-alleles 2' and '--max-alleles 2' arguments will filter out multi-allelic sites, as *Selscan* only accepts bi-allelic sites as input data to calculate metrics of extended haplotype homozygosity. As a final step, we also manually filtered out duplicate entries for the same variant in the resulting VCF file. We refer to this file as 'input_final.vcf' in the subsequent section ('*Selscan* (version 1.2.0)').

*Selscan* (version 1.2.0)

To calculate $nS_L$ values, we used the following command:

**./selscan --nsl --vcf input_final.vcf --maf 0.01 --threads 16 --out output**

Apart from the minor allele frequency cutoff value (0.01), we used the default settings for other parameters. We used 16 threads to carry out calculations of $nS_L$ via *Selscan*, as shown above. We used the maximum of the $nS_L$ values ($6^{th}$ column) recorded in the *.nsl.out file to represent each enhancer.

*Tabix* (version.0.2.6)

We used *Tabix* to compress the newly generated VCF files and create tab-delimited index (.tbi) files required for subsequent analyses via *PopGenome*. The commands used are as follows:

**./bgzip -c output.recode.vcf > output.vcf.gz**

**./tabix -p vcf output.vcf.gz**

*SLiM* (version 2.4.1) *and Simulation of Neutral Evolutions*

For each tissue, we used *SLiM* to simulate neutral evolution and generate VCF files of variants resulting from the action of neutral evolution alone, given past demographic events. We used *SLiM*'s implementation of Gravel et al's pre-computed parameters of human demographic history for our neutral simulations. More specifically, at the beginning of the simulation (i.e. generation 1), the ancestral African effective population size was set to 7,310, which next expanded to 14,474 approximately 148,000 years ago (i.e. 5,920 generations ago). Approximately 51,000 years ago (i.e. 2,040 generations ago), the non-Africans split from Africans; the initial effective population size of these non-Africans was set to 1,861. The migration rates between Africans and non-Africans were set to 15 x $10^{-5}$. Next, approximately 23,000 years ago (i.e. 920 generations ago), the above-mentioned non-African population split into European and East Asian populations, with the initial effective population size for East Asians set to 554. In the same generation, the European effective population size was reduced to 1,032. The following migration rates were established for the remainder of the simulation: 2.5 x $10^{-5}$ for between Africans and Europeans, 0.78 x $10^{-5}$ for between Africans and Asians, and 3.11 x $10^{-5}$ for between Europeans and East Asians. Between generations 57,080 and 58,000, the European and East Asian populations were set to experience increase in their effective population sizes: for Europeans, the exponential coefficient was 0.0038 and 0.0048 for East Asians. A fixed mutation rate of 2.36 x $10^{-8}$ and recombination rate of 1 x $10^{-8}$ were used. For each tissue, we defined the length of the genomic element being simulated as the length of average of the lengths of all autosomal enhancers expressed in a given tissue. After 58,000 generations (i.e. end of the simulation), we sampled 661, 503, and 504 individuals from the "simulated" African, European, and East Asian population, respectively, to match the number of individuals included in the 1000 Genomes Project dataset. We used the following arguments to ensure that pairs of genomes being sampled belonged to the same individual:

```
p1_sample = p1.individuals;
sampled_p1 = sample(p1_sample, 661);
p2_sample = p2.individuals;
```

```
sampled_p2 = sample(p2_sample, 503);
pe_sample = p3.individuals;
sampled_p3 = sample(p3_sample, 504);
```

p1, p2, and p3, correspond to the simulated African, European, and East Asian populations. Finally, to specify the output format as VCF, we used the arguments as shown below:

```
sampled_individuals = c(sampled_p1, sampled_p2, sampled_p3);
sampled_individuals.genomes.outputVCF();
```

Genomic regions spanning 50kb upstream and downstream of the average enhancer length of a given tissue were created using the same approach as described above, except that the recombination rate for each tissue was set to the values obtained as described in a previous section ("Calculation of Average Recombination Rates per Tissue"). For each tissue, we generated 1) 10,000 "neutrally simulated" VCF files that contain genotype information for the "simulated" enhancer region and 2) an additional 2,500 "neutrally simulated" VCF files that contain genotype information for the "simulated" region that spans 50kb upstream and downstream of a "simulated" enhancer. Calculation of Tajima's $D$, Weir & Cockerham's $F_{\mathrm{ST}}$, $nS_{\mathrm{L}}$, and H12 using these VCF files were carried out as described in previous sections. We refer to these metrics as values generated under "neutral expectations".

*GO Enrichment Analyses using TopGO (version 2.32.0)*

We first downloaded the GO annotations file (download date: May 17th, 2018) from:

http://www.geneontology.org/page/download-go-annotations/goa_human.gaf.gz

We processed the file downloaded from the above link and created a "Gene Universe (GU)" file for each tissue: this file would contain all the Gene Ontology (GO) IDs associated with any given putative target gene associated with an enhancer that is active in a given tissue. The GU file was created using only the autosomal putative target genes of enhancers in any given tissue. We also created "Target Genes (TG)" file for each tissue and each of the metrics we had calculated: more specifically, we obtained a list of GO IDs for the autosomal, putative target genes associated with enhancers that exhibit significantly extreme values of any given metric compared to those calculated on the neutrally simulated sequences. After further processing the files into a format required for *TopGO* analyses, we carried out the enrichment analyses via *TopGO* using the classic Fisher's test (statistic = "fisher") and the "weight" algorithm (algorithm = "weight"). This analysis was repeated for the three "root" GO terms ("Biological Process"; ontology = "BP", "Cellular Component"; ontology = "CC", "Molecular Function"; ontology = "MF"). We further carried out a FDR-based multiple test correction using R to obtain the final list of significantly enriched GO terms for each tissue of any given metric.

*Semantic Similarity Analyses using GOSemSim*

We used the results obtained from the preceding section ("GO Enrichment Analyses using TopGO") to calculate pairwise semantic similarity of the GO terms between tissues for the $nS_L$ metrics calculated in all three populations: this analysis was carried out only for the $nS_L$ metric, as only this metric had more than one tissue with at least 5 significantly enriched GO terms. We used the “org.Hs.eg.db” Bioconductor package for the genome-wide annotation data of humans, as follows:

**SemData <- godata(‘org.Hs.eg.db’, ont = “BP”, computeIC = “FALSE”)**

We changed the argument “BP” to “CC” and “MF” to run the calculations for the three different root GO terms. ‘computeIC’ argument was set to “FALSE” as we were using a graph-based method for our calculations. We carried out pairwise semantic similarity calculations using Wang’s graph-based method and combined the similarity scores via the Best-Match Average (BMA) strategy as follows:

**semantic_similarity <- mgoSim(first_tissue_data, second_tissue_data, SemData, measure = “Wang”, combine = “BMA”)**

‘first_tissue_data’ refers to a R object that contains the list of GO terms (obtained from the preceding section) for one tissue, while ‘second_tissue_data’ stores the list of GO terms for another tissue.

*Transposable Elements Analyses using BEDOPS (version 2.4.35)*

To obtain a genome-wide annotation of repeat regions, we downloaded the RepeatMasker (rmsk) track for the hg19 genome assembly (download date: June $15^{th}$, 2018) from UCSC genome browser:

http://hgdownload.cse.ucsc.edu/goldenpath/hg19/database/rmsk.txt.gz

We further converted this file into a BED format by grabbing the 6th, 7th, 8th, 12$^{th}$, 11$^{th}$, and 10$^{th}$ columns and carried out sorting using BEDOPS ("sort-bed"). We further filtered this file ("background_regions.bed") to include only those genomic regions that are annotated as the following: "LINE", "SINE", "LTR" ("background_TE_regions.bed"). We next took the enhancers that exhibit significant deviations from neutral expectations for each metric in each tissue and using the coordinates recorded in the original BED files, created respective BED files ("test_regions.bed"). We next used BEDOPS to overlap the two types of BED files as follows:

**bedmap --echo --echo-map-id-uniq test_regions.bed background_TE_regions.bed**

*GWAS catalog Data (v.1.0.2)*
The GWAS catalog was downloaded from (download date: June 2$^{nd}$, 2018):

https://www.ebi.ac.uk/gwas/docs/file-downloads/gwas_catalog_v1.0.2_associatons_final.gz

For any given tissue, we obtained a list of the rs IDs of the SNPs that lie within the enhancers that displayed statistically extreme values of the calculated metrics compared to those calculated on the neutrally simulated sequences, and used those SNPs to query the GWAS catalog and obtain a list of significantly associated traits.

*Linkage Disequilibrium Calculations using Plink (v1.90b6.2)*

We first processed the VCF files for each chromosome (downloaded as described in “1000 Genomes Project Phase 3 Genotype Dataset” Section) as follows: 1) we first filtered out indels (i.e., included only SNPs) and 2) included only Europeans (n = 503). To obtain a list of SNPs that are in linkage disequilibrium (i.e., r2 = 1) with those used in the preceding section (GWAS catalog Data (v.1.0.2), we calculated Linkage Disequilibrium (LD) using Plink as follows:

**./plink --vcf genotype_data.vcf --exclude duplicate_snps_list --ld-snp-list --r2 --inter-chr 1 --out output_prefix**

“genotype_data.vcf” refers to the filtered VCF file generated above and “duplicate_snps_list” is a file listing the duplicate rs IDs. We used the resulting SNPs’ rs IDs to query the GWAS catalog, as described in the preceding section.

**Figure S1. Distributions of metrics of recent positive selection in the 41 human tissues**

The Ridgeline plots depicting the distributions of the four different metrics calculated on all the autosomal enhancers within each tissue. (a) Tajima's $D$ (b) Weir & Cockerham's $F_{ST}$ (c) H12 (d) $nS_L$ (Africans) (e) $nS_L$ (Europeans) (f) $nS_L$ (East Asians) The dotted vertical lines indicate the 95th percentile values obtained from calculating the same metrics on the 10,000 (or 2,500 for the $nS_L$ metric) neutrally simulated sequences.

**Figure S2. Quantification of the semantic similarities of the functional enrichment among tissues**

The Heatmaps depicting the Semantic Similarity (SS) values calculated on the set of significantly enriched GO terms between any tissues for the $nS_L$ metrics. The SS values were calculated only for those tissues that have 5 or more significantly enriched GO terms. This analysis was done for all 3 populations, and all 3 root GO terms (Biological Process (BP), Molecular Functions (MF), Cellular Components (CC))

**Figure S3. Comparisons of the proportions of overlap with TE-annotated regions between enhancers with and without significant evidence of recent positive selection**

The Bar charts depicting the proportions of overlap with TE-annotated regions for enhancers with significant evidence of recent selection (teal) and without such evidence (gray). Bars with asterisks (*) above them indicate significant chi-square test results.

**Figure S4. Comparisons of the distributions of Tajima's *D* values between enhancers active in brain and testis and other tissues**

The Kernel Density Estimation (KDE) plots depicting the distributions of the Tajima's $D$ values in the brain and testis (pale violet red) and other tissues (gray) for the statistically significant pairwise comparisons.

**Figure S5. Comparisons of the distributions of Weir & Cockerham's $F_{ST}$ values between enhancers active in brain and testis and other tissues**

The Kernel Density Estimation (KDE) plots depicting the distributions of the Weir & Cockerham's $F_{ST}$ values in the brain and testis (pale violet red) and other tissues (gray) for the statistically significant pairwise comparisons.

**Figure S6. Comparisons of the distributions of H12 values between enhancers active in brain and other tissues**

The Kernel Density Estimation (KDE) plots depicting the distributions of the H12 values in the brain (pale violet red) and other tissues (gray) for the statistically significant pairwise comparisons.

**Figure S7. Comparisons of the distributions of H12 values between enhancers active in testis and other tissues**

The Kernel Density Estimation (KDE) plots depicting the distributions of the H12 values in the testis (pale violet red) and other tissues (gray) for the statistically significant pairwise comparisons.

**Figure S8. Comparisons of median values of the calculated metrics across tissues**

Tissues have been ranked from highest to lowest (or lowest from highest for Tajima's *D*) median values of the calculated metrics. The colors of the bars correlate with the number of enhancers included in the tissues. Brain and testis have been bolded in the x-axis to ease comparison with other tissues.

**Figure S9. Comparisons of proportions of enhancers with significant evidence of recent positive selection across tissues**

Tissues have been ranked from highest to lowest proportions of enhancers with significant evidence of recent positive selection for each metric. The colors of the bars correlate with the number of enhancers included in the tissues. Brain and testis have been bolded in the x-axis to ease comparison with other tissues.

**Figure S10. Comparisons of the distributions of Tajima's *D* values between tissue-specific and tissue-broad enhancers**

The violin plots depicting the distributions of the values of Tajima's $D$ for the tissue-specific (light purple) and tissue-broad (light green) enhancers for all 41 tissues. Plots with asterisks (*) above them indicate significant pairwise comparisons.

**Figure S11. Comparisons of the distributions of Weir & Cockerham's $F_{ST}$ values between tissue-specific and tissue-broad enhancers**

The violin plots depicting the distributions of the values of Weir & Cockerham's $F_{ST}$ for the tissue-specific (light purple) and tissue-broad (light green) enhancers for all 41 tissues. Plots with asterisks (*) above them indicate significant pairwise comparisons.

**Figure S12. Comparisons of the distributions of H12 values between tissue-specific and tissue-broad enhancers**

The violin plots depicting the distributions of the values of H12 for the tissue-specific (light purple) and tissue-broad (light green) enhancers for all 41 tissues. Plots with asterisks (*) above them indicate significant pairwise comparisons.

**Figure S13. Comparisons of the distributions $nS_L$ (Africans) values between tissue-specific and tissue-broad enhancers**

The violin plots depicting the distributions of the values of $nS_L$ (Africans) for the tissue-specific (light purple) and tissue-broad (light green) enhancers for all 41 tissues. Plots with asterisks (*) above them indicate significant pairwise comparisons.

**Figure S14. Comparisons of the distributions $nS_L$ (Europeans) values between tissue-specific and tissue-broad enhancers**

The violin plots depicting the distributions of the values of $nS_L$ (Europeans) for the tissue-specific (light purple) and tissue-broad (light green) enhancers for all 41 tissues. Plots with asterisks (*) above them indicate significant pairwise comparisons.

**Figure S15. Comparisons of the distributions $nS_L$ (East Asians) values between tissue-specific and tissue-broad enhancers**

The violin plots depicting the distributions of the values of $nS_L$ (East Asians) for the tissue-specific (light purple) and tissue-broad (light green) enhancers for all 41 tissues. Plots with asterisks (*) above them indicate significant pairwise comparisons.

Figure S1

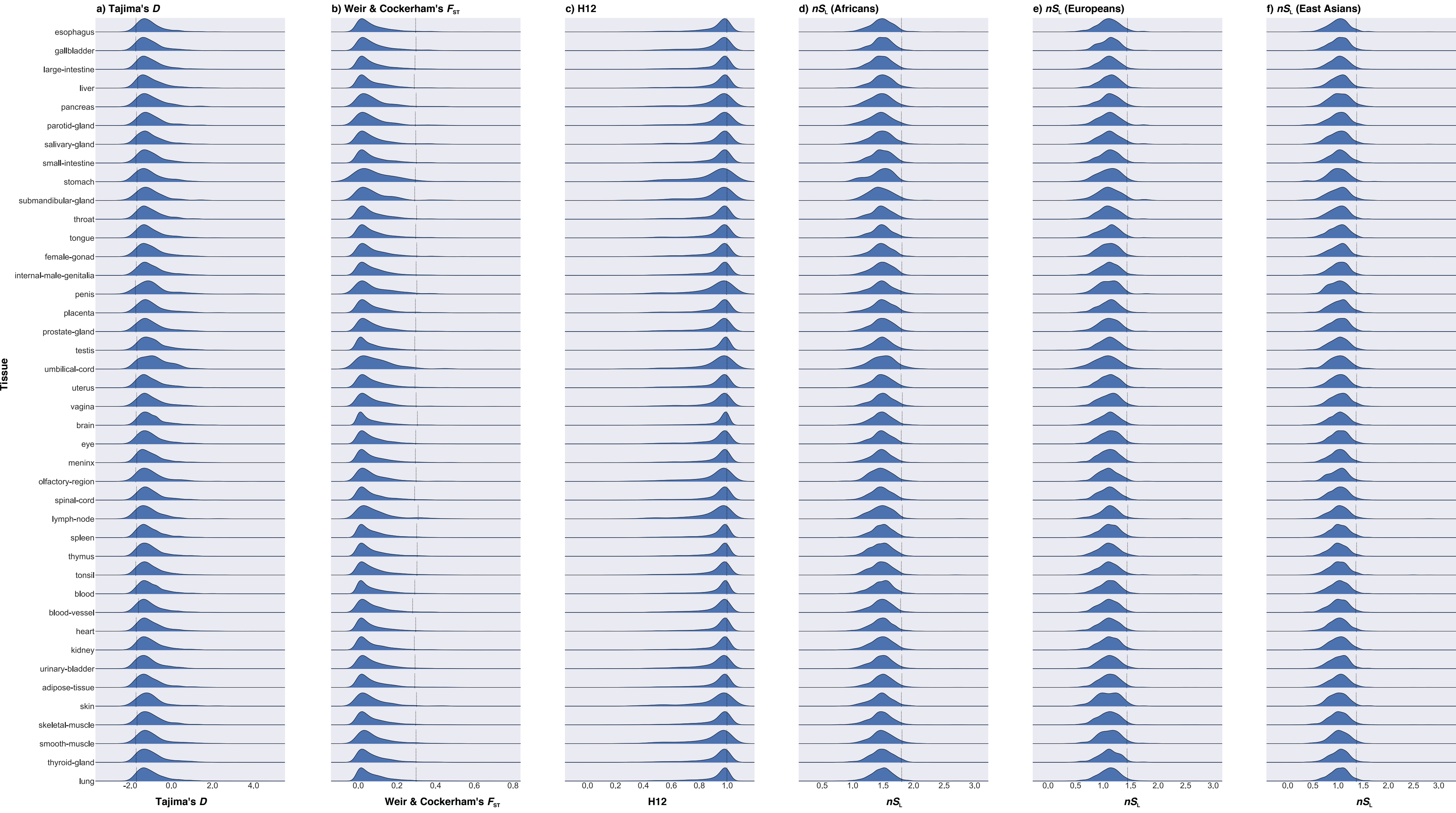
a) Tajima's D
b) Weir & Cockerham's $F_{ST}$
c) H12
d) $nS_L$ (Africans)
e) $nS_L$ (Europeans)
f) $nS_L$ (East Asians)
Tissue
esophagus
gallbladder
large-intestine
liver
pancreas
parotid-gland
salivary-gland
small-intestine
stomach
submandibular-gland
throat
tongue
female-gonad
internal-male-genitalia
penis
placenta
prostate-gland
testis
umbilical-cord
uterus
vagina
brain
eye
meninx
olfactory-region
spinal-cord
lymph-node
spleen
thymus
tonsil
blood
blood-vessel
heart
kidney
urinary-bladder
adipose-tissue
skin
skeletal-muscle
smooth-muscle
thyroid-gland
lung
-2.0 0.0 2.0 4.0
Tajima's D
0.0 0.2 0.4 0.6 0.8
Weir & Cockerham's $F_{ST}$
0.0 0.2 0.4 0.6 0.8 1.0
H12
0.5 1.0 1.5 2.0 2.5 3.0
$nS_L$
0.0 0.5 1.0 1.5 2.0 2.5 3.0
$nS_L$
0.0 0.5 1.0 1.5 2.0 2.5 3.0
$nS_L$

Figure S2

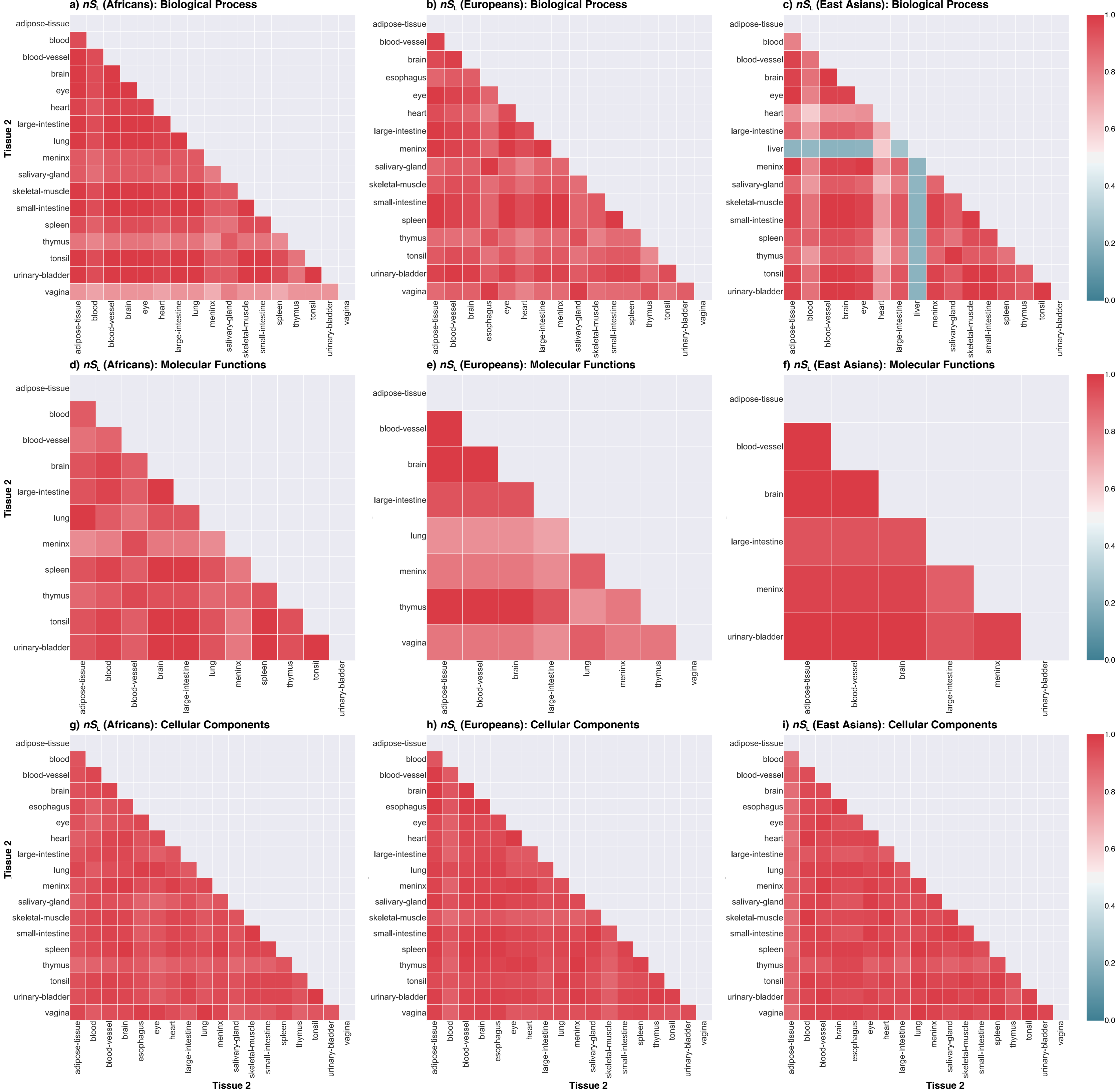
a) $nS_L$ (Africans): Biological Process
b) $nS_L$ (Europeans): Biological Process
c) $nS_L$ (East Asians): Biological Process
d) $nS_L$ (Africans): Molecular Functions
e) $nS_L$ (Europeans): Molecular Functions
f) $nS_L$ (East Asians): Molecular Functions
g) $nS_L$ (Africans): Cellular Components
h) $nS_L$ (Europeans): Cellular Components
i) $nS_L$ (East Asians): Cellular Components
Tissue 2
adipose-tissue
blood
blood-vessel
brain
esophagus
eye
heart
large-intestine
liver
lung
meninx
salivary-gland
skeletal-muscle
small-intestine
spleen
thymus
tonsil
urinary-bladder
vagina
0.0
0.2
0.4
0.6
0.8
1.0

Figure S3

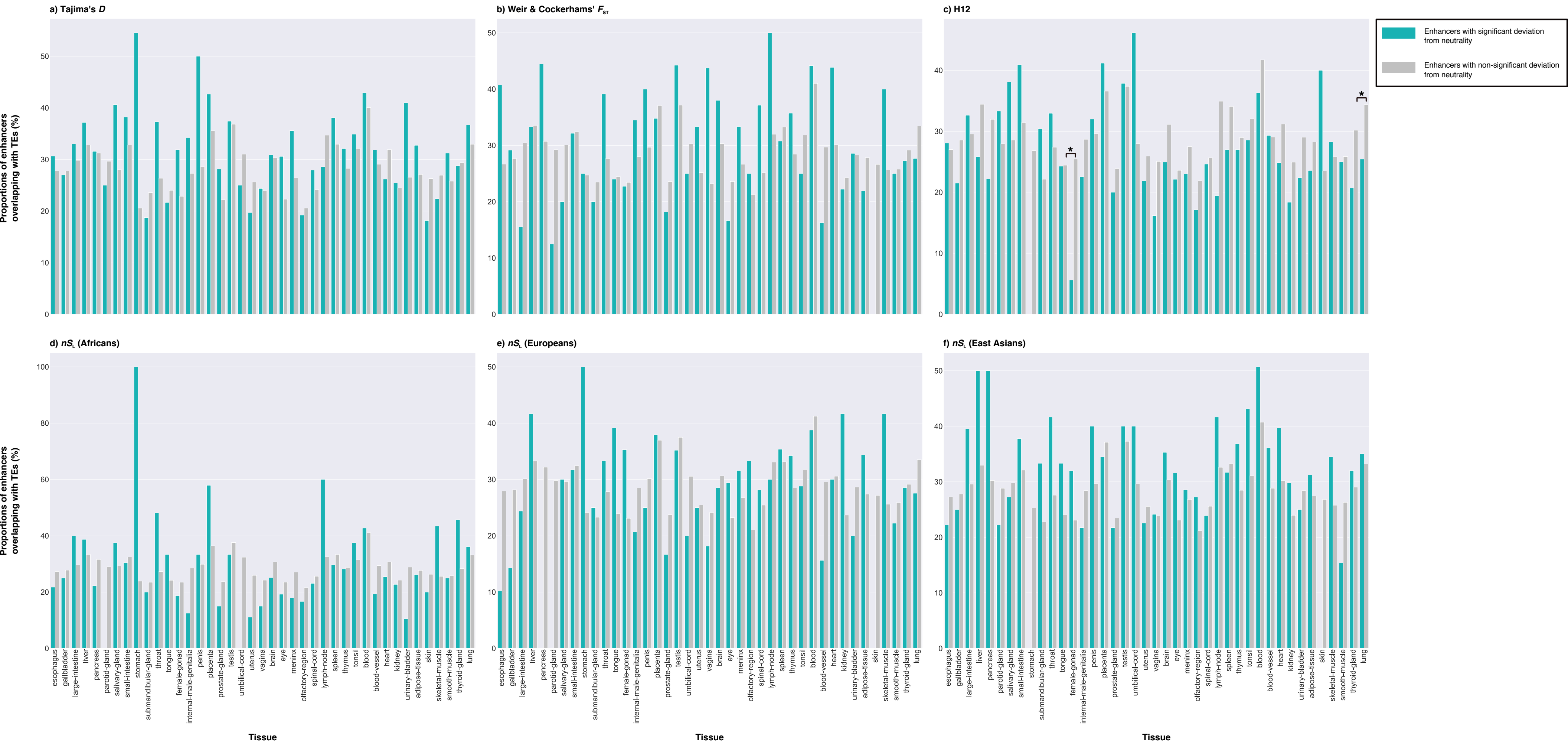

a) Tajima's D
b) Weir & Cockerhams' $F_{ST}$
c) H12
d) $nS_L$ (Africans)
e) $nS_L$ (Europeans)
f) $nS_L$ (East Asians)
Enhancers with significant deviation from neutrality
Enhancers with non-significant deviation from neutrality
Proportions of enhancers overlapping with TEs (%)
Tissue
esophagus
gallbladder
large-intestine
liver
pancreas
parotid-gland
salivary-gland
small-intestine
stomach
submandibular-gland
throat
tongue
female-gonad
internal-male-genitalia
penis
placenta
prostate-gland
testis
umbilical-cord
uterus
vagina
brain
eye
menix
olfactory-region
spinal-cord
lymph-node
spleen
thymus
tonsil
blood
blood-vessel
heart
kidney
urinary-bladder
adipose-tissue
skin
skeletal-muscle
smooth-muscle
thyroid-gland
lung

Figure S4

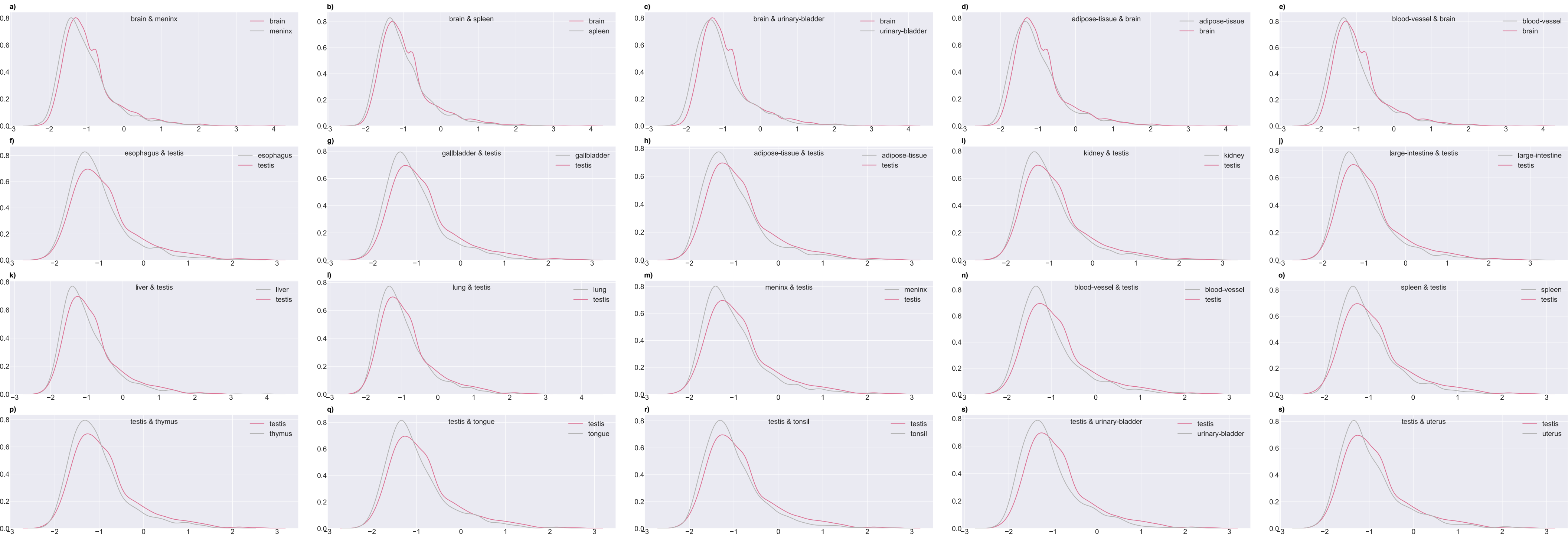

a) brain & meninx
brain
meninx
b) brain & spleen
brain
spleen
c) brain & urinary-bladder
brain
urinary-bladder
d) adipose-tissue & brain
adipose-tissue
brain
e) blood-vessel & brain
blood-vessel
brain
f) esophagus & testis
esophagus
testis
g) gallbladder & testis
gallbladder
testis
h) adipose-tissue & testis
adipose-tissue
testis
i) kidney & testis
kidney
testis
j) large-intestine & testis
large-intestine
testis
k) liver & testis
liver
testis
l) lung & testis
lung
testis
m) meninx & testis
meninx
testis
n) blood-vessel & testis
blood-vessel
testis
o) spleen & testis
spleen
testis
p) testis & thymus
testis
thymus
q) testis & tongue
testis
tongue
r) testis & tonsil
testis
tonsil
s) testis & urinary-bladder
testis
urinary-bladder
s) testis & uterus
testis
uterus

Figure S5

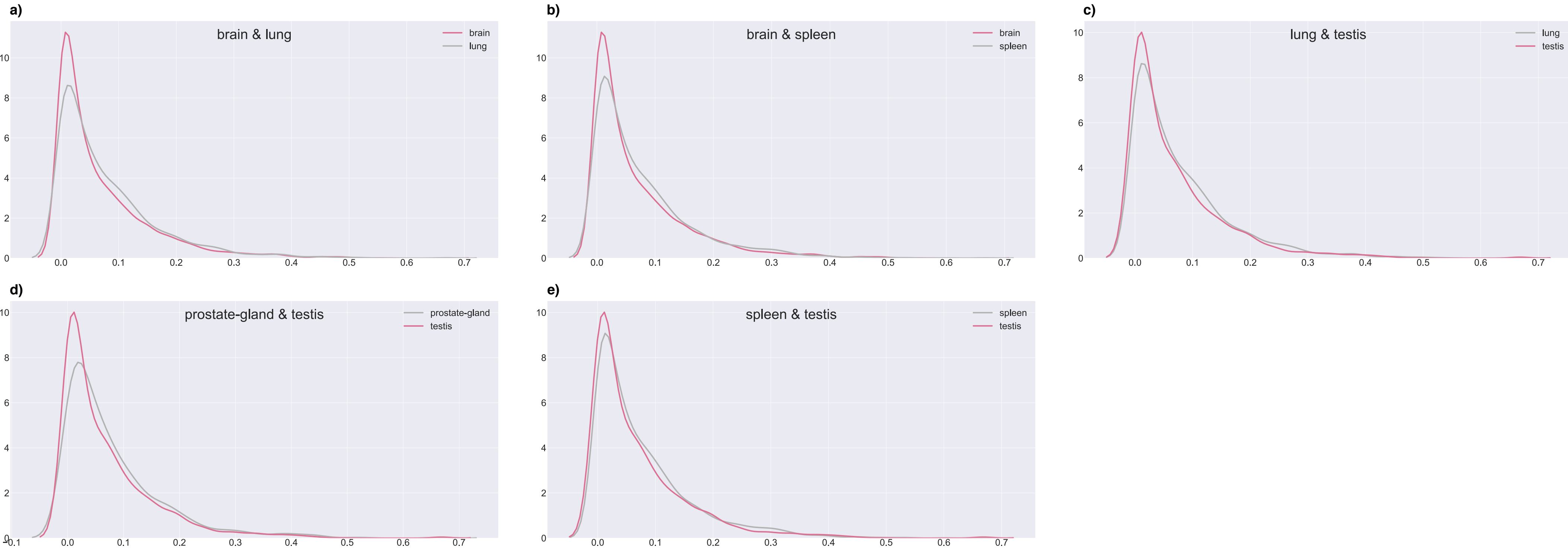
a)
brain & lung
brain
lung
b)
brain & spleen
brain
spleen
c)
lung & testis
lung
testis
d)
prostate-gland & testis
prostate-gland
testis
e)
spleen & testis
spleen
testis

Figure S6

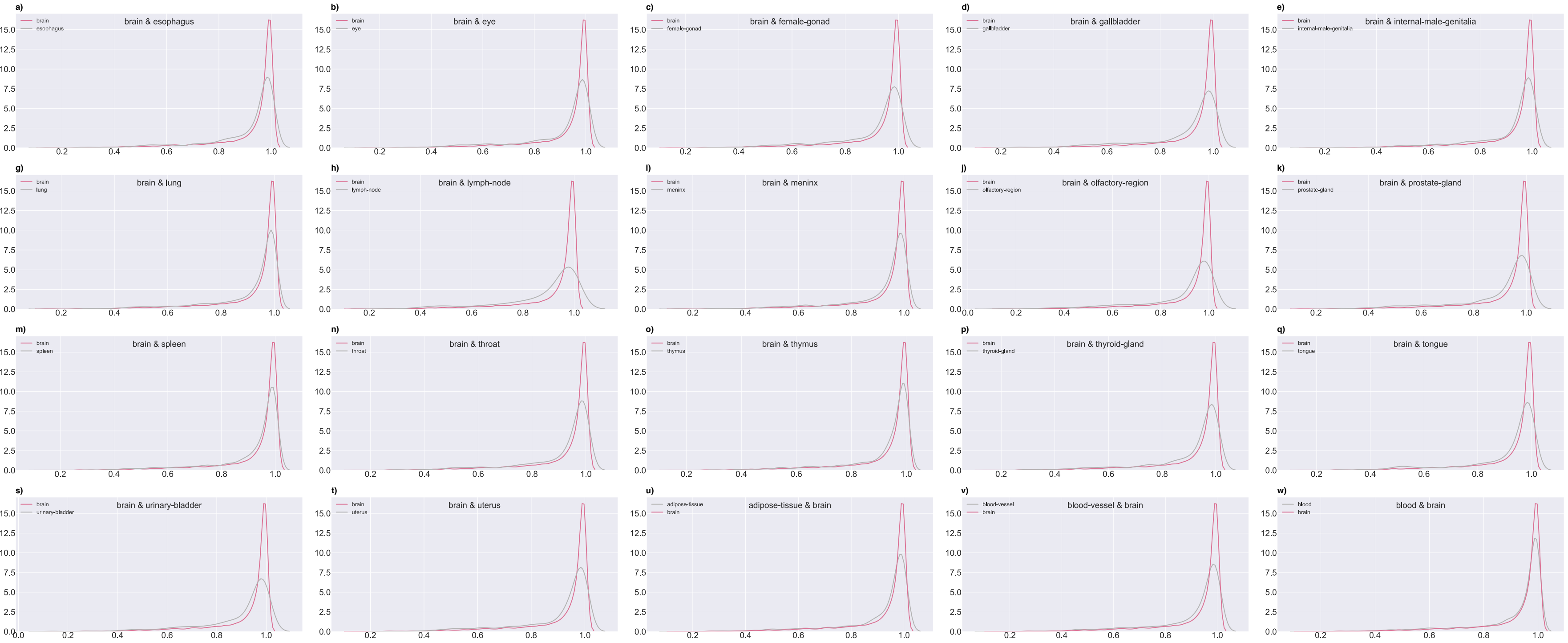

a) brain & esophagus
brain
esophagus
b) brain & eye
brain
eye
c) brain & female-gonad
brain
female-gonad
d) brain & gallbladder
brain
gallbladder
e) brain & internal-male-genitalia
brain
internal-male-genitalia
f) brain & large-intestine
brain
large-intestine
g) brain & lung
brain
lung
h) brain & lymph-node
brain
lymph-node
i) brain & meninx
brain
meninx
j) brain & olfactory-region
brain
olfactory-region
k) brain & prostate-gland
brain
prostate-gland
l) brain & smooth-muscle
brain
smooth-muscle
m) brain & spleen
brain
spleen
n) brain & throat
brain
throat
o) brain & thymus
brain
thymus
p) brain & thyroid-gland
brain
thyroid-gland
q) brain & tongue
brain
tongue
r) brain & tonsil
brain
tonsil
s) brain & urinary-bladder
brain
urinary-bladder
t) brain & uterus
brain
uterus
u) adipose-tissue & brain
adipose-tissue
brain
v) blood-vessel & brain
blood-vessel
brain
w) blood & brain
blood
brain

Figure S7

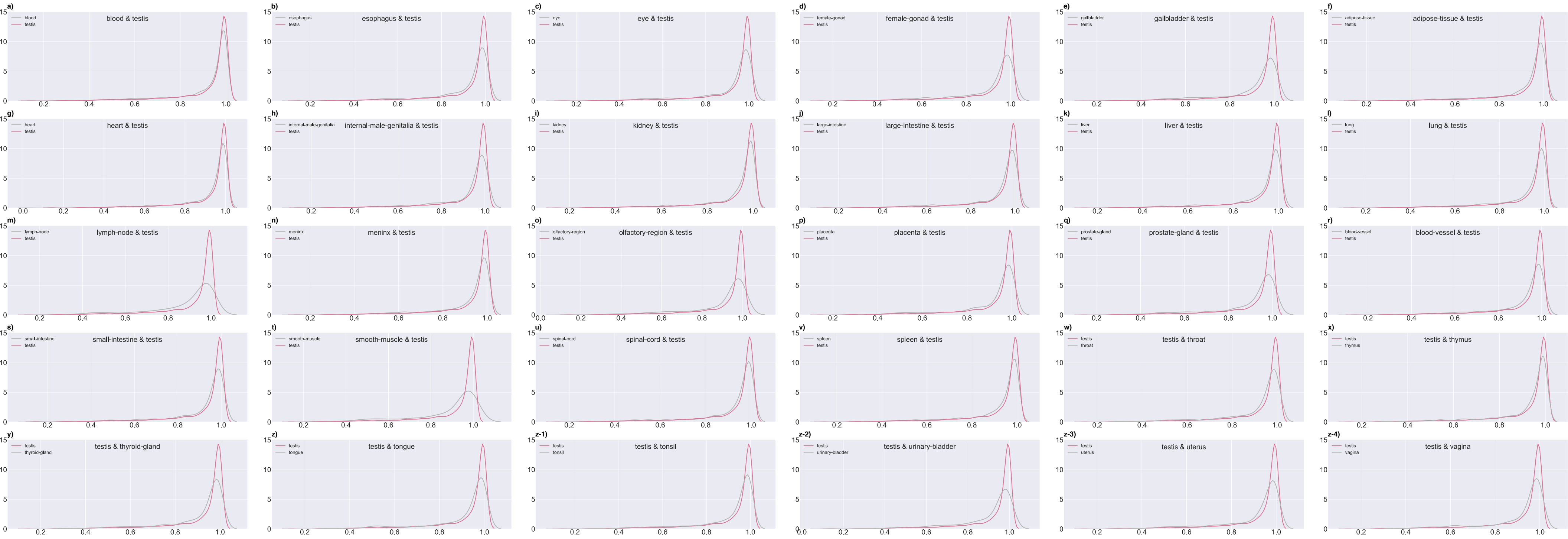

a) blood & testis
b) esophagus & testis
c) eye & testis
d) female-gonad & testis
e) gallbladder & testis
f) adipose-tissue & testis
g) heart & testis
h) internal-male-genitalia & testis
i) kidney & testis
j) large-intestine & testis
k) liver & testis
l) lung & testis
m) lymph-node & testis
n) meninx & testis
o) olfactory-region & testis
p) placenta & testis
q) prostate-gland & testis
r) blood-vessel & testis
s) small-intestine & testis
t) smooth-muscle & testis
u) spinal-cord & testis
v) spleen & testis
w) testis & throat
x) testis & thymus
y) testis & thyroid-gland
z) testis & tongue
z-1) testis & tonsil
z-2) testis & urinary-bladder
z-3) testis & uterus
z-4) testis & vagina

Figure S8

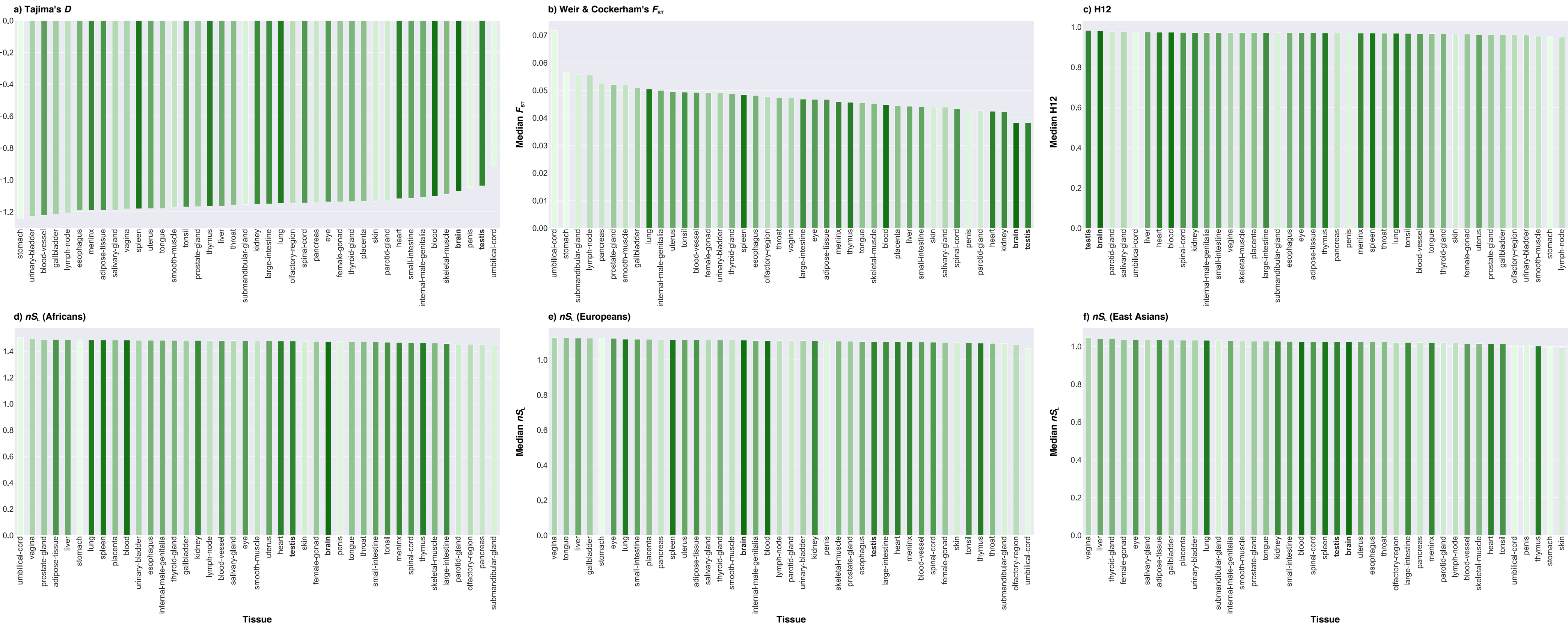

a) Tajima's D
Median Tajima's D
b) Weir & Cockerham's F_ST
Median F_ST
c) H12
Median H12
d) nS_L (Africans)
Median nS_L
e) nS_L (Europeans)
Median nS_L
f) nS_L (East Asians)
Median nS_L
Tissue
Tissue
Tissue

### Figure S9

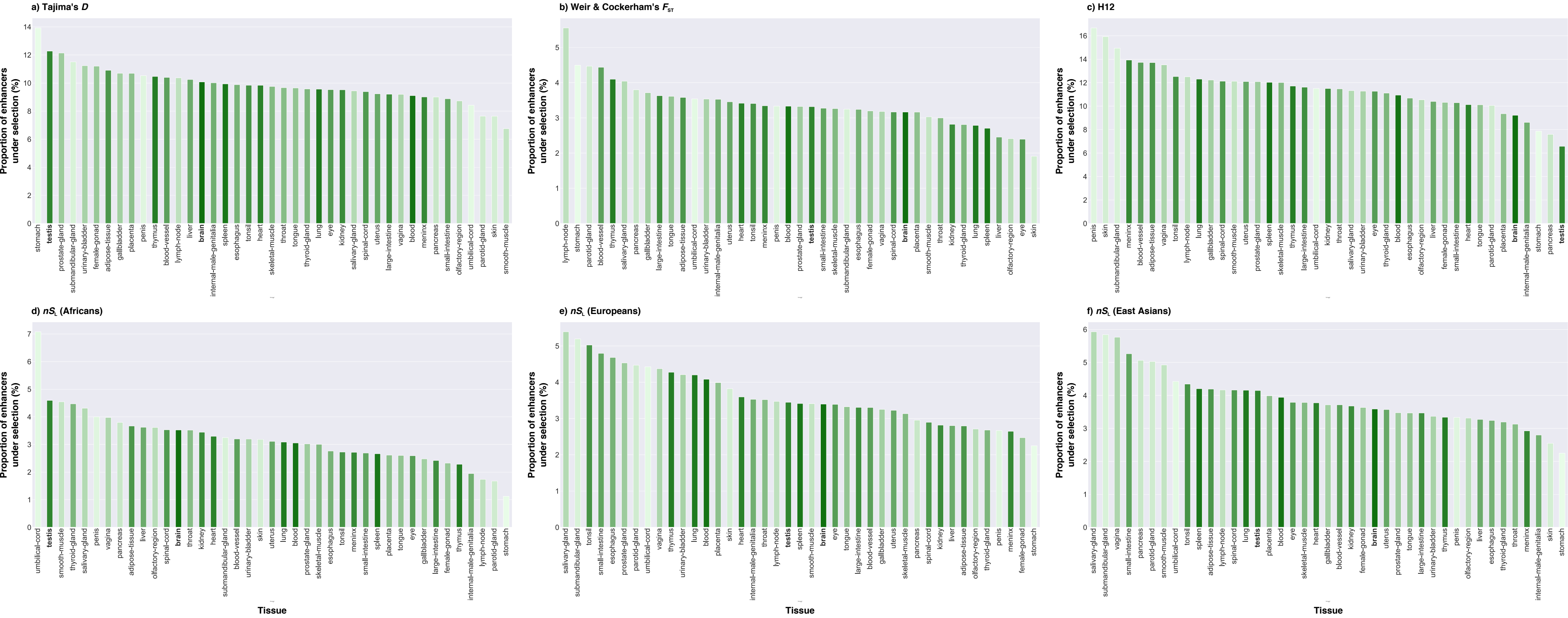
a) Tajima's D
Proportion of enhancers under selection (%)
stomach testis prostate-gland submandibular-gland urinary-bladder female-gonad adipose-tissue gallbladder placenta penis thymus blood-vessel lymph-node liver brain internal-male-genitalia spleen esophagus tonsil heart skeletal-muscle throat tongue thyroid-gland lung eye kidney salivary-gland spinal-cord uterus large-intestine vagina blood meninx pancreas small-intestine olfactory-region umbilical-cord parotid-gland skin smooth-muscle
b) Weir & Cockerham's $F_{ST}$
Proportion of enhancers under selection (%)
lymph-node stomach parotid-gland blood-vessel thymus salivary-gland pancreas gallbladder large-intestine tongue adipose-tissue umbilical-cord urinary-bladder internal-male-genitalia uterus heart tonsil meninx penis blood prostate-gland testis small-intestine skeletal-muscle submandibular-gland esophagus female-gonad vagina spinal-cord brain placenta smooth-muscle throat kidney thyroid-gland lung spleen liver olfactory-region eye skin
c) H12
Proportion of enhancers under selection (%)
penis skin submandibular-gland meninx blood-vessel adipose-tissue vagina tonsil lymph-node lung gallbladder spinal-cord smooth-muscle uterus prostate-gland spleen skeletal-muscle thymus large-intestine umbilical-cord kidney throat salivary-gland urinary-bladder eye thyroid-gland blood esophagus olfactory-region liver female-gonad small-intestine heart tongue parotid-gland placenta brain internal-male-genitalia stomach pancreas testis
d) $nS_L$ (Africans)
Proportion of enhancers under selection (%)
umbilical-cord testis smooth-muscle thyroid-gland salivary-gland penis vagina pancreas adipose-tissue liver olfactory-region spinal-cord brain throat kidney heart submandibular-gland blood-vessel urinary-bladder skin uterus lung blood prostate-gland skeletal-muscle esophagus tonsil meninx small-intestine spleen placenta tongue eye gallbladder large-intestine female-gonad thymus internal-male-genitalia lymph-node parotid-gland stomach
Tissue
e) $nS_L$ (Europeans)
Proportion of enhancers under selection (%)
salivary-gland submandibular-gland tonsil small-intestine esophagus prostate-gland parotid-gland umbilical-cord vagina thymus urinary-bladder lung blood placenta skin heart internal-male-genitalia throat lymph-node testis spleen smooth-muscle brain eye tongue large-intestine blood-vessel gallbladder uterus skeletal-muscle pancreas spinal-cord kidney liver adipose-tissue olfactory-region thyroid-gland penis meninx female-gonad stomach
Tissue
f) $nS_L$ (East Asians)
Proportion of enhancers under selection (%)
salivary-gland submandibular-gland vagina small-intestine pancreas parotid-gland smooth-muscle umbilical-cord tonsil spleen adipose-tissue lymph-node spinal-cord lung testis placenta blood eye skeletal-muscle heart gallbladder blood-vessel kidney female-gonad brain uterus prostate-gland tongue large-intestine urinary-bladder thymus penis olfactory-region liver esophagus thyroid-gland throat meninx internal-male-genitalia skin stomach
Tissue

Figure S10

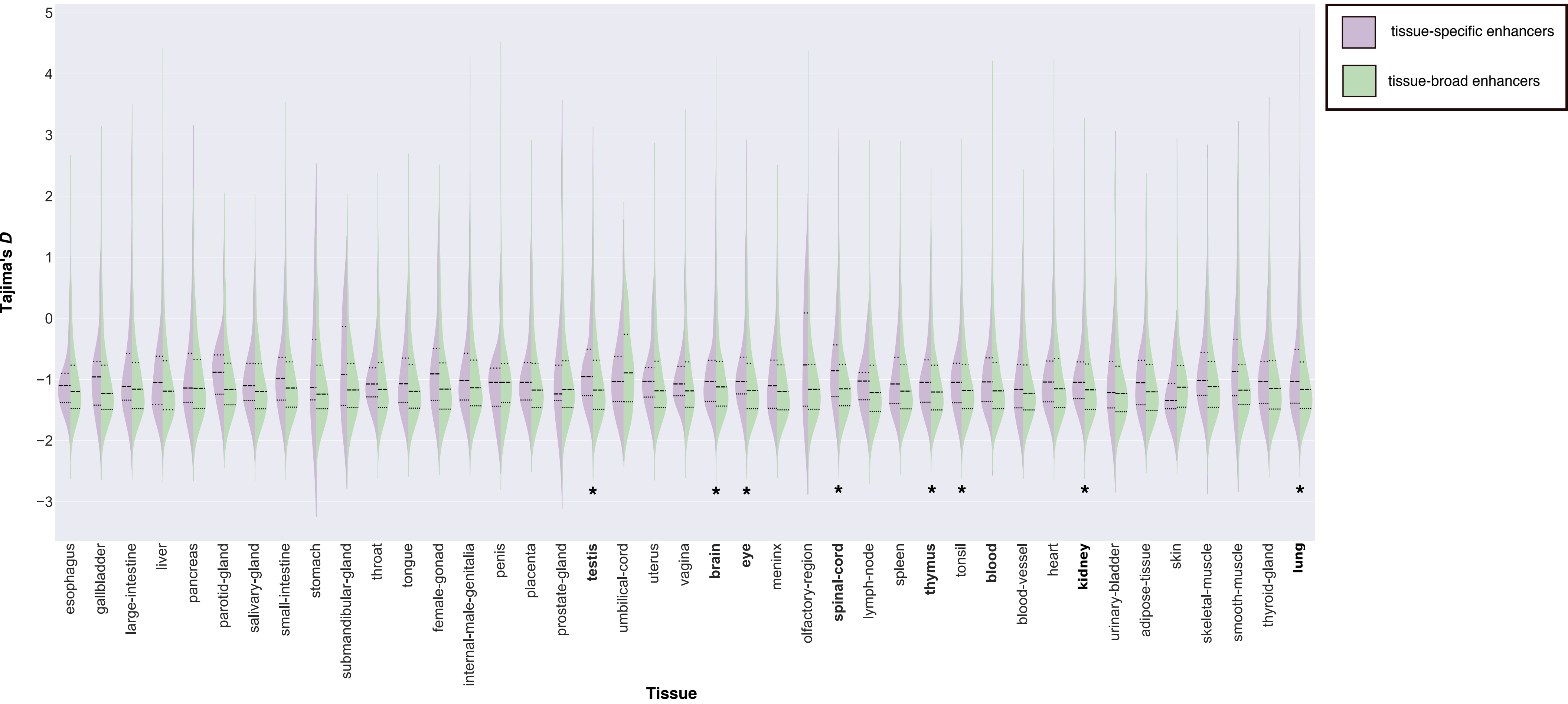

tissue-specific enhancers
tissue-broad enhancers
Tajima's D
Tissue
esophagus
gallbladder
large-intestine
liver
pancreas
parotid-gland
salivary-gland
small-intestine
stomach
submandibular-gland
throat
tongue
female-gonad
internal-male-genitalia
penis
placenta
prostate-gland
testis
umbilical-cord
uterus
vagina
brain
eye
meninx
olfactory-region
spinal-cord
lymph-node
spleen
thymus
tonsil
blood
blood-vessel
heart
kidney
urinary-bladder
adipose-tissue
skin
skeletal-muscle
smooth-muscle
thyroid-gland
lung

Figure S11

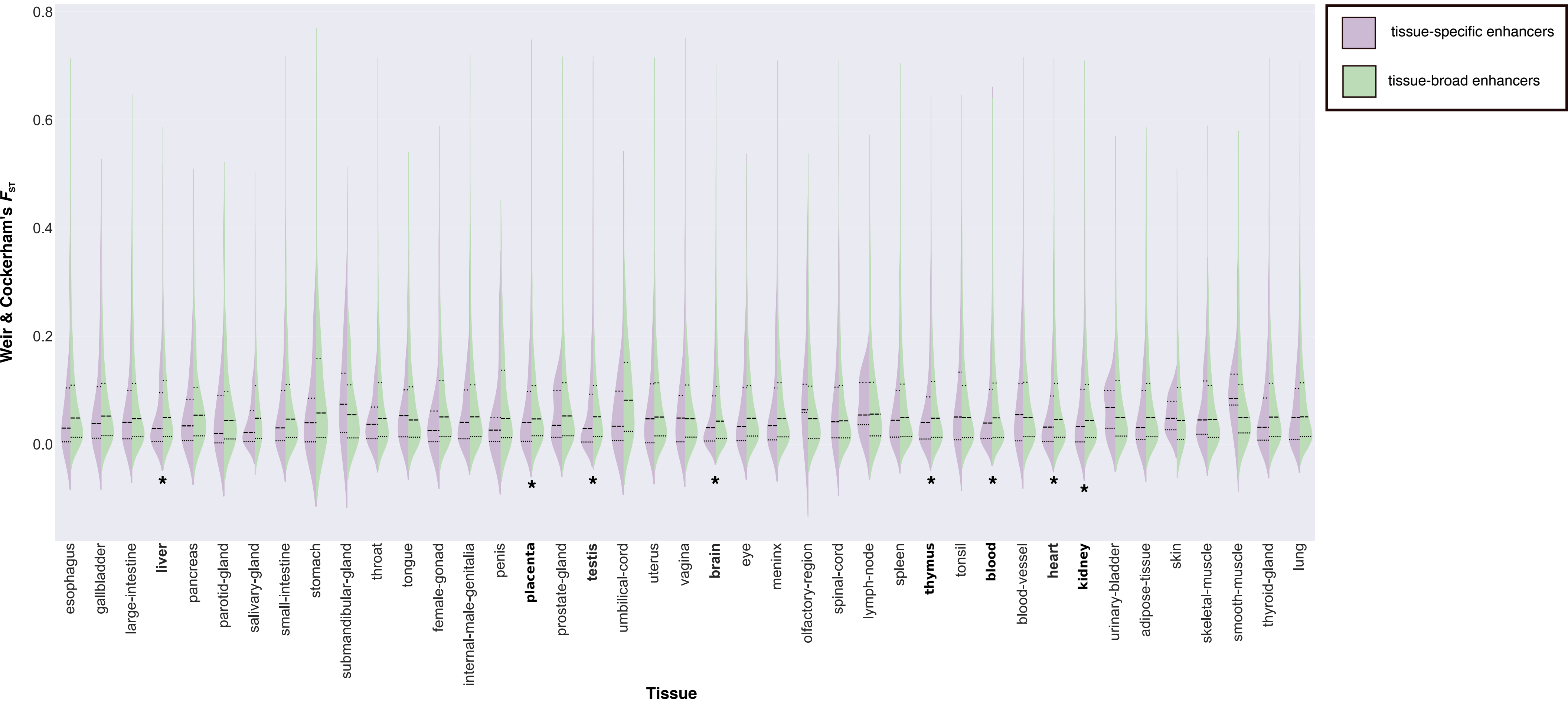

0.8
0.6
0.4
0.2
0.0
Weir & Cockerham's $F_{ST}$
tissue-specific enhancers
tissue-broad enhancers
esophagus
gallbladder
large-intestine
liver
pancreas
parotid-gland
salivary-gland
small-intestine
stomach
submandibular-gland
throat
tongue
female-gonad
internal-male-genitalia
penis
placenta
prostate-gland
testis
umbilical-cord
uterus
vagina
brain
eye
meninx
olfactory-region
spinal-cord
lymph-node
spleen
thymus
tonsil
blood
blood-vessel
heart
kidney
urinary-bladder
adipose-tissue
skin
skeletal-muscle
smooth-muscle
thyroid-gland
lung
Tissue

### Figure S12

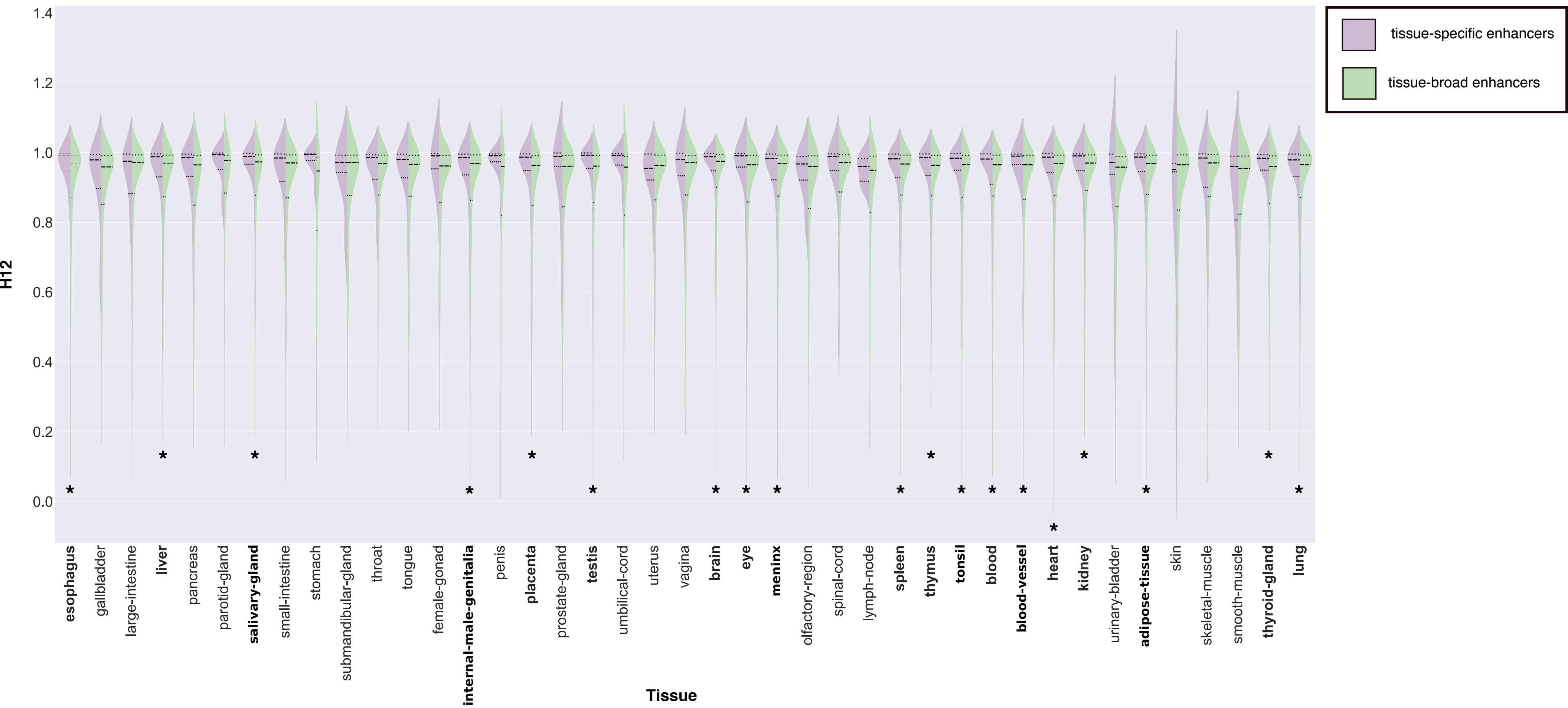
H12
Tissue
tissue-specific enhancers
tissue-broad enhancers
1.4
1.2
1.0
0.8
0.6
0.4
0.2
0.0
esophagus
gallbladder
large-intestine
liver
pancreas
parotid-gland
salivary-gland
small-intestine
stomach
submandibular-gland
throat
tongue
female-gonad
internal-male-genitalia
penis
placenta
prostate-gland
testis
umbilical-cord
uterus
vagina
brain
eye
meninx
olfactory-region
spinal-cord
lymph-node
spleen
thymus
tonsil
blood
blood-vessel
heart
kidney
urinary-bladder
adipose-tissue
skin
skeletal-muscle
smooth-muscle
thyroid-gland
lung

Figure S13

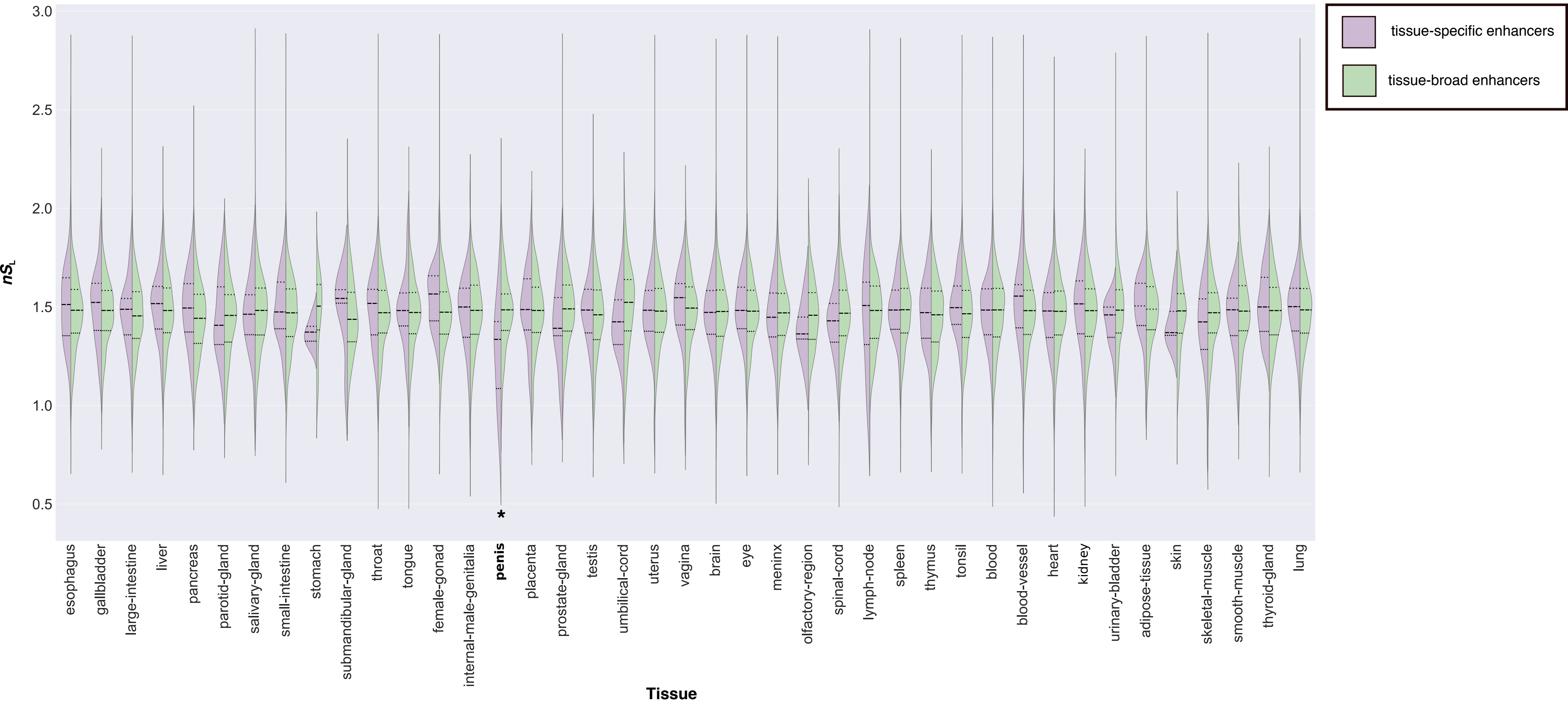

3.0
2.5
2.0
1.5
1.0
0.5
$nS_L$
*
esophagus
gallbladder
large-intestine
liver
pancreas
parotid-gland
salivary-gland
small-intestine
stomach
submandibular-gland
throat
tongue
female-gonad
internal-male-genitalia
penis
placenta
prostate-gland
testis
umbilical-cord
uterus
vagina
brain
eye
meninx
olfactory-region
spinal-cord
lymph-node
spleen
thymus
tonsil
blood
blood-vessel
heart
kidney
urinary-bladder
adipose-tissue
skin
skeletal-muscle
smooth-muscle
thyroid-gland
lung
Tissue
tissue-specific enhancers
tissue-broad enhancers

Figure S14

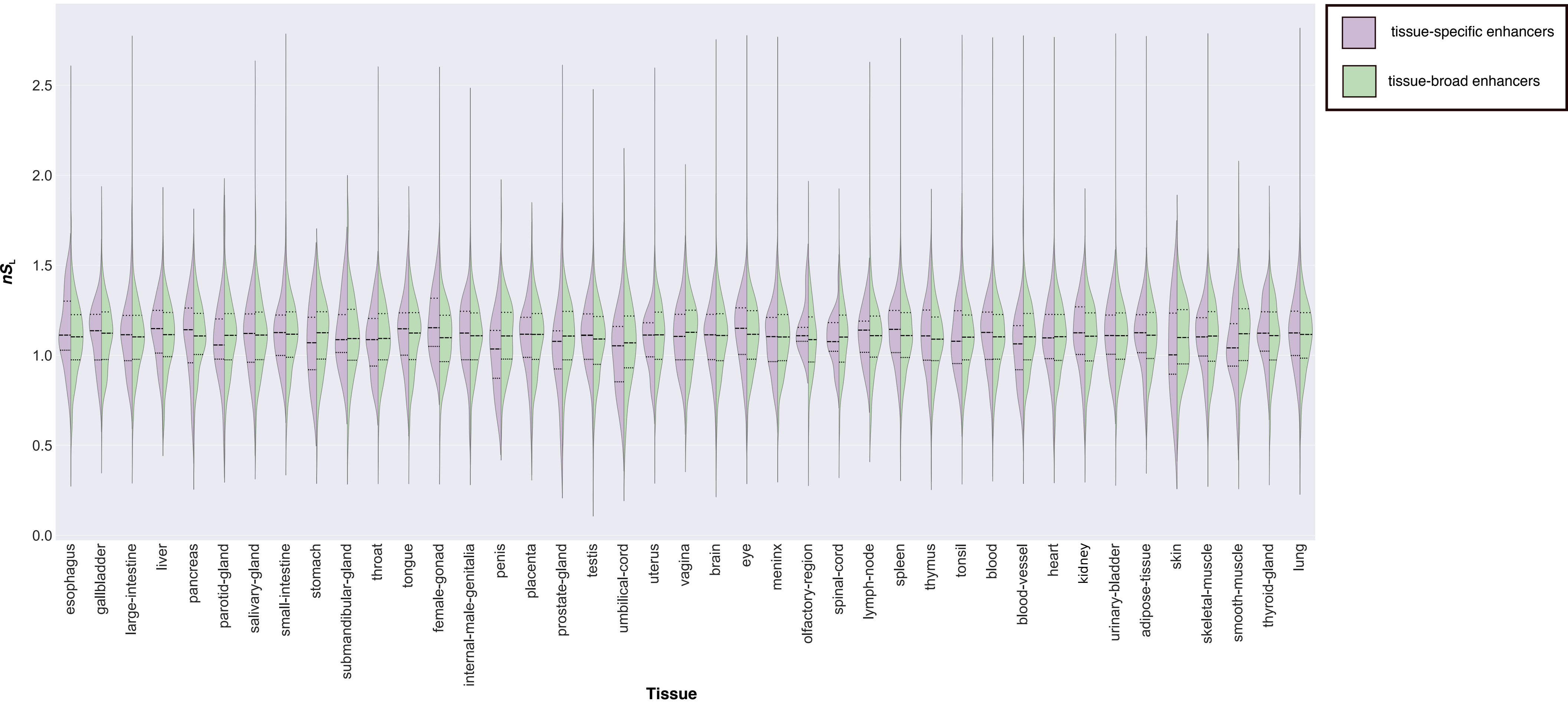

tissue-specific enhancers
tissue-broad enhancers
$nS_L$
0.0
0.5
1.0
1.5
2.0
2.5
esophagus
gallbladder
large-intestine
liver
pancreas
parotid-gland
salivary-gland
small-intestine
stomach
submandibular-gland
throat
tongue
female-gonad
internal-male-genitalia
penis
placenta
prostate-gland
testis
umbilical-cord
uterus
vagina
brain
eye
meninx
olfactory-region
spinal-cord
lymph-node
spleen
thymus
tonsil
blood
blood-vessel
heart
kidney
urinary-bladder
adipose-tissue
skin
skeletal-muscle
smooth-muscle
thyroid-gland
lung
Tissue

### Figure S15

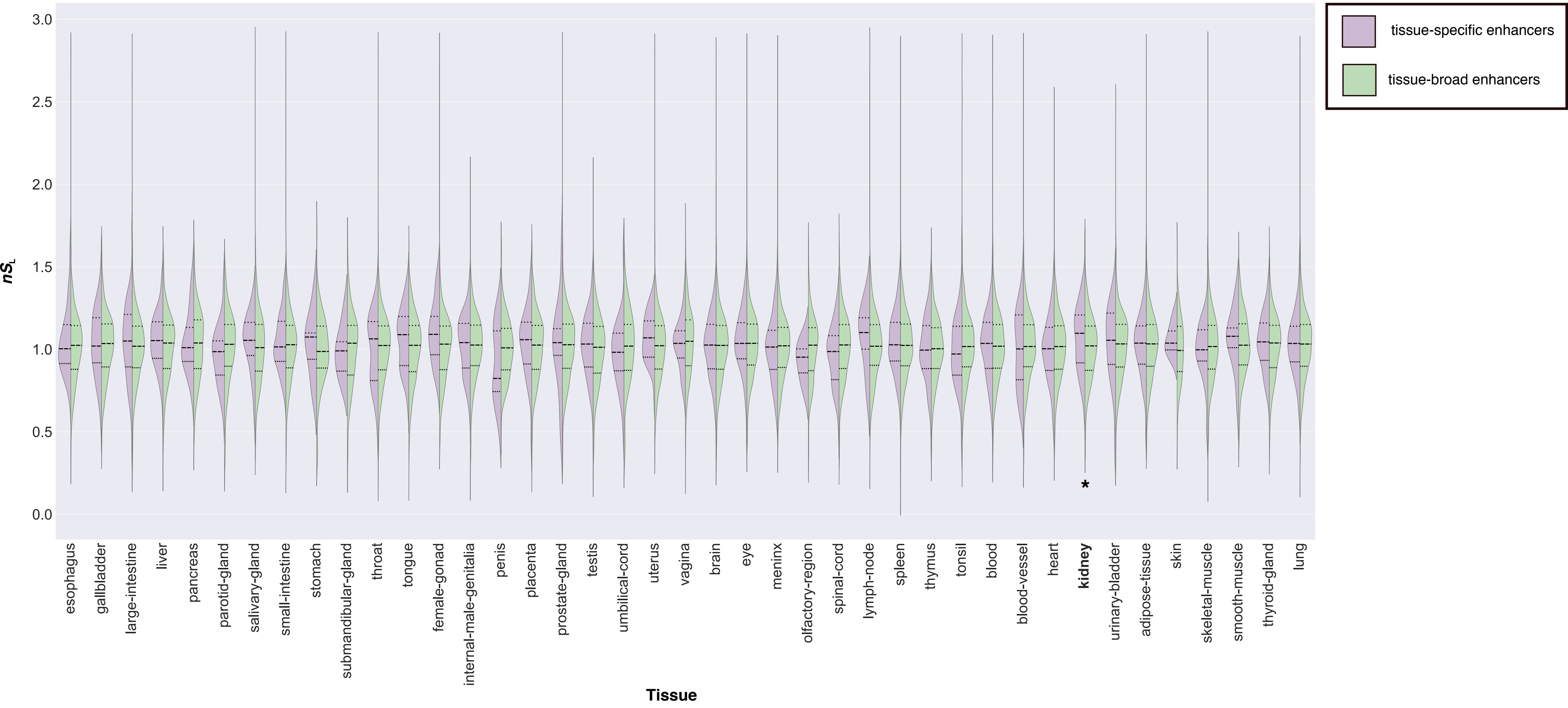

tissue-specific enhancers
tissue-broad enhancers
$nS_L$
3.0
2.5
2.0
1.5
1.0
0.5
0.0
esophagus
gallbladder
large-intestine
liver
pancreas
parotid-gland
salivary-gland
small-intestine
stomach
submandibular-gland
throat
tongue
female-gonad
internal-male-genitalia
penis
placenta
prostate-gland
testis
umbilical-cord
uterus
vagina
brain
eye
meninx
olfactory-region
spinal-cord
lymph-node
spleen
thymus
tonsil
blood
blood-vessel
heart
kidney
*
urinary-bladder
adipose-tissue
skin
skeletal-muscle
smooth-muscle
thyroid-gland
lung
Tissue

**Table S1. Number of enhancers included in each tissue**

| Tissue | Number of enhancers |
|---|---|
| **Esophagus** | 845 |
| **Gallbladder** | 654 |
| **Large-intestine** | 1,266 |
| **Liver** | 875 |
| **Pancreas** | 243 |
| **Parotid-gland** | 184 |
| **Salivary-gland** | 387 |
| **Small-intestine** | 880 |
| **Stomach** | 92 |
| **Submandibular-gland** | 159 |
| **Throat** | 782 |
| **Tongue** | 708 |
| **Female-gonad** | 693 |
| **Internal-male-genitalia** | 846 |
| **Penis** | 154 |
| **Placenta** | 735 |
| **Prostate-gland** | 667 |
| **Testis** | 1,621 |
| **Umbilical-cord** | 115 |
| **Uterus** | 877 |
| **Vagina** | 512 |
| **Brain** | 4,883 |
| **Eye** | 1,019 |
| **Meninx** | 1,447 |
| **Olfactory-region** | 338 |
| **Spinal-cord** | 1,115 |
| **Lymph-node** | 292 |
| **Spleen** | 2,429 |

| | |
|---|---|
| **Thymus** | 1,741 |
| **Tonsil** | 1,193 |
| **Blood** | 3,657 |
| **Blood-vessel** | 982 |
| **Heart** | 1,737 |
| **Kidney** | 1,294 |
| **Urinary-bladder** | 602 |
| **Adipose-tissue** | 1,161 |
| **Skin** | 161 |
| **Skeletal-muscle** | 784 |
| **Smooth-muscle** | 270 |
| **Thyroid-gland** | 795 |
| **Lung** | 2,366 |

**Table S2. Proportions of enhancers with significant evidence of recent positive selection according to each metric within a given tissue**

| Tissue | $F_{ST}$ | Tajima's $D$ | H12 | $nS_L$ (Africans) | $nS_L$ (Europeans) | $nS_L$ (East Asians) |
|---|---|---|---|---|---|---|
| Adipose-tissue | 3.581 | 10.907 | 13.712 | 3.668 | 2.795 | 4.192 |
| Blood-vessel | 4.438 | 10.4 | 13.725 | 3.199 | 3.302 | 3.715 |
| Blood | 3.331 | 9.105 | 10.938 | 3.054 | 4.081 | 3.942 |
| Brain | 3.165 | 10.077 | 9.221 | 3.524 | 3.397 | 3.587 |
| Esophagus | 3.241 | 9.881 | 10.684 | 2.761 | 4.682 | 3.241 |
| Eye | 2.393 | 9.529 | 11.266 | 2.592 | 3.390 | 3.789 |
| Female-gonad | 3.198 | 11.201 | 10.320 | 2.326 | 2.471 | 3.634 |
| Gallbladder | 3.715 | 10.696 | 12.229 | 2.477 | 3.251 | 3.715 |
| Heart | 3.415 | 9.837 | 10.126 | 3.295 | 3.595 | 3.775 |
| Internal-male-genitalia | 3.528 | 10.014 | 8.617 | 1.946 | 3.528 | 2.798 |
| Kidney | 2.817 | 9.516 | 11.502 | 3.443 | 2.817 | 3.678 |
| large-intestine | 3.629 | 9.205 | 11.613 | 2.419 | 3.306 | 3.468 |
| Liver | 2.453 | 10.250 | 10.397 | 3.621 | 2.804 | 3.271 |
| Lung | 2.786 | 9.563 | 12.302 | 3.086 | 4.201 | 4.158 |
| Lymph-node | 5.556 | 10.370 | 12.5 | 1.736 | 3.472 | 4.167 |
| Meninx | 3.343 | 9.008 | 13.928 | 2.716 | 2.646 | 2.925 |
| Olfactory-region | 2.410 | 8.725 | 10.542 | 3.614 | 2.711 | 3.313 |
| Pancreas | 3.797 | 9.005 | 7.595 | 3.797 | 2.954 | 5.063 |
| Parotid-gland | 4.469 | 7.643 | 10.056 | 1.676 | 4.469 | 5.028 |
| Penis | 3.333 | 10.526 | 16.667 | 4 | 2.667 | 3.333 |
| Placenta | 3.164 | 10.692 | 9.354 | 2.613 | 3.989 | 3.989 |
| Prostate-gland | 3.323 | 12.137 | 12.085 | 3.021 | 4.532 | 3.474 |
| Salivary-gland | 4.043 | 9.440 | 11.321 | 4.313 | 5.391 | 5.930 |
| Skeletal-muscle | 3.264 | 9.753 | 12.010 | 3.003 | 3.133 | 3.786 |
| Skin | 1.911 | 7.639 | 15.924 | 3.185 | 3.822 | 2.548 |
| Small-intestine | 3.275 | 8.877 | 10.292 | 2.690 | 4.795 | 5.263 |
| Smooth-muscle | 3.030 | 6.751 | 12.121 | 4.545 | 3.409 | 4.924 |
| Spinal-cord | 3.167 | 9.384 | 12.127 | 3.529 | 2.896 | 4.163 |
| Spleen | 2.705 | 9.936 | 12.027 | 2.663 | 3.412 | 4.203 |
| Stomach | 4.494 | 13.924 | 7.865 | 1.124 | 2.247 | 2.247 |
| Submandibular-gland | 3.247 | 11.511 | 14.935 | 3.247 | 5.195 | 5.844 |
| Testis | 3.318 | 12.274 | 6.573 | 4.595 | 3.446 | 4.148 |

| | | | | | | |
|---|---|---|---|---|---|---|
| Throat | 2.999 | 9.668 | 11.473 | 3.520 | 3.520 | 3.129 |
| Thymus | 4.098 | 10.465 | 11.710 | 2.283 | 4.274 | 3.337 |
| Thyroid-gland | 2.810 | 9.579 | 11.111 | 4.470 | 2.682 | 3.193 |
| Tongue | 3.613 | 9.646 | 10.116 | 2.601 | 3.324 | 3.468 |
| Tonsil | 3.407 | 9.842 | 12.521 | 2.726 | 5.026 | 4.344 |
| Umbilical-cord | 3.540 | 8.421 | 11.504 | 7.080 | 4.425 | 4.425 |
| Urinary-bladder | 3.535 | 11.234 | 11.279 | 3.199 | 4.209 | 3.367 |
| Uterus | 3.456 | 9.233 | 12.097 | 3.111 | 3.226 | 3.571 |
| Vagina | 3.181 | 9.193 | 13.519 | 3.976 | 4.374 | 5.765 |

All values have been rounded to the 3$^{rd}$ decimal place.
